# Disulfide Crosslinking Induces Rapid Degradation of Arc/Arg3.1 via Hsp70-Mediated Ubiquitin Ligase Pathway

**DOI:** 10.1101/2025.07.20.665809

**Authors:** Dami So, In-Kang Song, Yeon Jung Kim, Yeonjoo Lee, Yeon Seung Park, Hee-Jung Kim, Kong-Joo Lee, Eun Joo Song

## Abstract

Activity-regulated cytoskeleton-associated protein (Arc/Arg3.1) is an immediate-early gene (IEG) induced by stress and synaptic activity, characterized by transient expression and rapid degradation. However, the mechanisms governing its degradation remain unclear. In this study, we identify a novel degradation pathway for Arc/Arg3.1, driven by its structural features. We demonstrate that the proteasomal degradation of Arc/Arg3.1 is modulated by the Hsp70-CHIP complex, with ubiquitination being impaired in HSF1 knockout cells. The formation of a Cys34-Cys159 disulfide bond crosslinks Arc/Arg3.1 into high-molecular-weight oligomers, altering its ubiquitination pattern and degradation kinetics compared to the C159A mutant. Hydrogen-deuterium exchange mass spectrometry (HDX-MS) revealed that wild-type (WT) Arc/Arg3.1 adopts a more compact structure than the C159A mutant. Notably, the C159A mutant fails to interact with HSF1, resulting in Hsp70 induction upon heat shock. Our findings propose a feedback loop in which disulfide crosslinking of Arc/Arg3.1 induces rapid degradation through Hsp70-mediated ubiquitination, which in turn modulates the heat shock response by inhibiting HSF1 function.

## 1. Introduction

Activity-regulated cytoskeleton-associated protein (Arc/Arg3.1) is an attractive marker of neuronal activity, known for its crucial roles in learning and memory through the regulation of neuronal plasticity, including long-term potentiation (LTP), long-term depression (LTD), and homeostatic plasticity (1). Arc/Arg3.1 interacts with dynamin, a large GTPase essential for intracellular membrane trafficking including clathrin-mediated synaptic vesicle recycling, and endophilin, a protein involved in vesicle formation and function (2–5). The expression level of Arc/Arg3.1 is tightly regulated. Arc/Arg3.1, as an immediate-early gene (IEG), is expressed at low levels under resting status, but its transcription is rapidly and transiently induced in response to external stimuli including brain-derived neurotrophic factor (BDNF) activation, metabotropic glutamate receptor (mGluR) activation and so on (6–9). Interestingly, Arc/Arg3.1 is markedly induced by heat shock and other cellular stress inducers including diamide, sodium arsenite and H_2_O_2_, in the neuronal and non-neuronal systems (10).

Arc/Arg3.1 not only responds to stress but also acts as a negative regulator of the heat shock response (HSR) (10). Specifically, Arc/Arg3.1 inhibits the binding of Heat Shock Factor 1 (HSF1) to heat shock elements (HSEs) in gene promoters, thereby reducing the transcription of Hsp70 and Hsp27 at both mRNA and protein levels. HSF1, the principal transcription factor in HSR, is activated in response to stress through phosphorylation, trimerization, and nuclear translocation, where it initiates the transcription of heat shock genes (11, 12). Active HSF1 trimers are subsequently inactivated by interactions with Hsp70, Hsp40, or heterogeneous nuclear ribonucleoprotein K (hnRNP K) (13), which diminishes their DNA-binding capability. By interfering with HSF1 function, Arc/Arg3.1 plays a critical role in modulating the cellular response to stress.

The degradation of Arc/Arg3.1 is also an essential aspect of its regulation, particularly in the context of its role in synaptic plasticity and stress responses. In neuronal systems, Arc/Arg3.1 is ubiquitinated by several E3 ubiquitin ligases, including Triad3A/RNF216 (14), E6AP/UBE3A (15) and the carboxy-terminus of Hsp70-interacting protein (CHIP) containing three tetratricopeptide repeat domains (16), resulting in its rapid proteasomal degradation. CHIP is known to regulate the ubiquitination and degradation of RIPK3, PTEN, HIF1A, TRAF2/6, and Smad1/Smad4 to regulate multiple signal transductions (17–24), and also interacts with Tau, Parkin, α-Synuclein, LRRK2, Ataxin1, and Ataxin3, which are involved in the pathogenesis of various neurodegenerative diseases (25–30). As a result, mutations in Triad3A/RNF216 and CHIP are linked to defective Arc/Arg3.1 ubiquitination in neuronal disorders such Gordon Holmes syndrome (GHS) (31) and spinocerebellar autosomal recessive 16 (SCAR16) (16). It was recently discovered that the autophagy/lysosomal pathway is responsible for at least some of the degradation of Arc/Arg3.1(32).

Despite the wealth of knowledge regarding Arc/Arg3.1’s role in the neuronal system, the mechanisms underlying its regulation and degradation in other cellular contexts, particularly under stress conditions, remain less well understood. Recent studies have highlighted that structural changes, such as aggregation or oxidation of proteins, can prompt their recognition and ubiquitination by E3 ligases, leading to degradation via the UPS (33, 34). Arc/Arg3.1, which contains a retroviral/retrotransposon GAG-like domain, is capable of self-association to form oligomers and virus-like capsids that transport its own mRNA within neuronal cells (35, 36). This self-association is thought to be mediated by the N-terminal helical coil, a motif critical for oligomerization (37).

Since Arc/Arg3.1 is a negative regulator of HSF1, through its inhibition of the HSF1 association to HSE in response to heat shock (10), we explored the heat shock response in HSF1-/- cells and found that the degradation of Arc/Arg3.1 is blocked in the absence of HSF1. This inhibition is abolished by reintroducing Hsp70 to HSF1-/- cells. This indicates that Hsp70 is required for the degradation of Arc/Arg3.1 via a negative feedback loop. CHIP, the E3-ligase associated with Hsp70, is known to promote the degradation of Arc/Arg3.1 (16). In this study, we examined the degradation of Arc/Arg3.1 using various mutants of CHIP and found that the complex of CHIP and Hsp70 is required to induce Arc/Arg3.1 degradation.

Furthermore, we explored the role of Arc/Arg3.1 oligomerization in its degradation process. Our study demonstrates that Arc/Arg3.1 readily oligomerizes under non-reducing conditions through disulfide crosslinking between Cys34 and Cys159, which was confirmed by cysteine mutant experiments and nanoUPLC-ESI-q-TOF tandem MS employing DBond algorithm (38) and structural modeling. The Arc/Arg3.1 C159A mutant is not oligomerized and not readily degraded by the UPS. The structural changes in this mutant were identified by hydrogen/deuterium exchange mass spectrometry (HDX-MS). The C159A mutant lost contact between the N-terminal domain (NTD) and the C-terminal domain (CTD), which is required for oligomerization (39). These findings suggest that Cys34-Cys159 disulfide crosslinking is a pre-requisite for the rapid degradation of Arc/Arg3.1 and regulates the function following structural changes by altering the interacting proteins. The results suggest that Arc/Arg3.1 degradation is finely regulated by structural changes with disulfide crosslinking and a negative feedback loop between Hsp70 and Arc/Arg3.1 with molecular interaction.

## 2. Results

### 2.1. UPS-dependent Arc/Arg3.1 degradation is inhibited in HSF1-/- cells

Arc/Arg3.1 is transiently expressed in response to heat shock-like stresses and readily degraded by the UPS in non-neuronal system (10). To determine the degradation mechanism of Arc/Arg3.1 in the feedback loop of the HSR, we examined the effect of HSF1 on Arc/Arg3.1 expression. Wild type (WT) and HSF1-/- mouse embryonic fibroblast (MEF) cells were exposed to heat shock (45°C for 15 min) and recovered in fresh media at 37°C for the indicated time. Arc/Arg3.1 was transiently induced and then disappeared in WT cells, while Arc/Arg3.1 was induced and accumulated during recovery in HSF1-/- cells (Fig. 1A and 1B). Arc/Arg3.1 protein is known to be readily degraded through the UPS, similar to other IEGs. To determine whether the accumulation of Arc/Arg3.1 in HSF1-/- cells was caused by a blockage of the UPS, we examined the Arc/Arg3.1 expression in WT cells treated with the proteasome inhibitor, MG132, and compared it with those in HSF1-/- cells. In response to heat shock, induced Arc/Arg3.1 proteins did not disappear in WT cells treated with MG132 during recovery. The accumulated levels of Arc/Arg3.1 in WT cells with MG132 were similar to those of HSF1-/- cells without MG132 (Fig. 1C and 1D). The results indicate that the UPS involved in Arc/Arg3.1 degradation is blocked in the HSF1-/- cells. This is further supported by the fact that the addition of MG132 did not alter Arc/Arg3.1 expression in HSF1-/- cells.

**Fig. 1.**
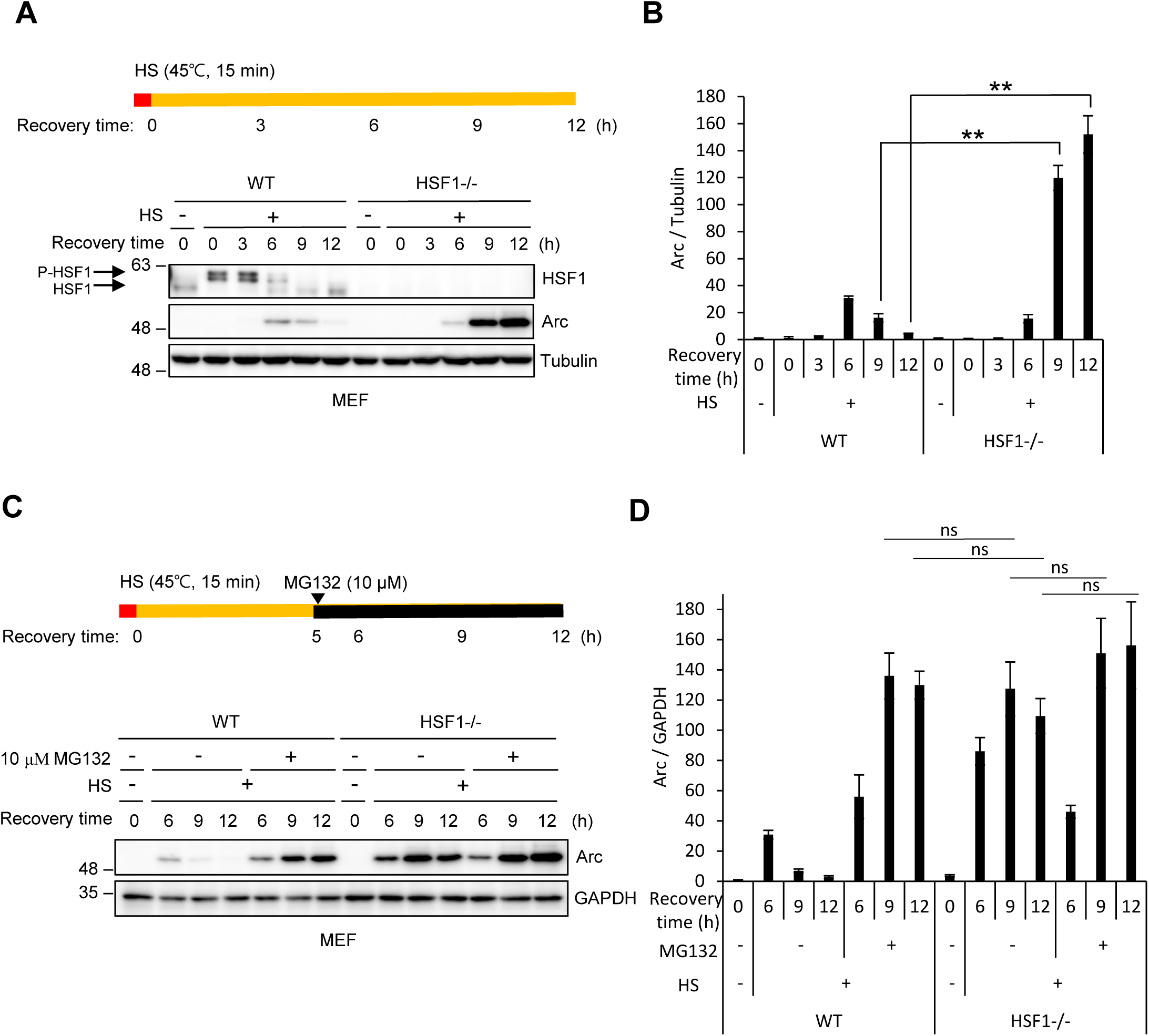
UPS-dependent Arc/Arg3.1 degradation is blocked in HSF1-/- cells. (A, B) Heat shock factor 1 (HSF1) knockout-/- cells accumulate Arc/Arg3.1 protein. WT and HSF1-/- MEF cells were exposed to heat shock at 45 °C for 15 min and allowed to recover in fresh media at 37 °C for the indicated times. HSF1 and Arc/Arg3.1 were detected by Western blot analysis with their specific antibodies. As a loading control, the tubulin level was detected by an anti-tubulin antibody. Western blot results were selected as representative data from triplicated experiments. Quantified results of triplicated Western blot images are shown in (B). (C, D) UPS-dependent Arc/Arg3.1 degradation is blocked in HSF1-/- cells. WT and HSF1-/- MEF cells were exposed to heat shock at 45 °C for 15 min and allowed to recover at 37° C for 5 h. Then, cells were incubated with 10 μM MG132 for the remaining indicated times. Arc/Arg3.1 were detected by Western blot analysis with anti-Arc antibodies. As a loading control, the GAPDH level was detected with an anti-GAPDH antibody. Western blot results were selected as representative data from triplicated experiments. Quantified results of triplicated Western images are shown in (D). Data are presented as the mean ± S.D. of triplicated experiments (t-test; **P < 0.01; NS denotes not significant).

### 2.2. Ectopic Hsp70 expression compensates for the effect of HSF1-/- in inhibiting the degradation of Arc/Arg3.1 and facilitates the degradation of Arc/Arg3.1

Since HSF1 does not affect the expression and activity of the proteasome (40) and HSF1-/- cells accumulate short-lived ubiquitinated proteins (41), we focused on chaperone-assisted UPS or degradation. To investigate whether accumulation of Arc/Arg3.1 protein in HSF1-/- cells is due to the absence of specific Hsps, we examined the expression of chaperones. HSF1-/- cells do not express Hsp70, Hsp40 and Hsp27 among multiple chaperones (Fig. 2A) when cells were exposed to heat shock and recovered for the indicated time. Because CHIP has been previously shown to ubiquitinate Arc/Arg3.1(16) and because Hsp70 controls bound substrate degradation by ubiquitination in conjunction with CHIP (42), we postulated that Arc/Arg3.1 degradation aided by Hsp70 is inhibited in HSF1-/- cells. To investigate whether the reduction in inducible Hsp70 expression in HSF1-/- cells contributes to inhibiting the degradation of Arc/Arg3.1, HSF1-/- cells transiently transfected with Flag empty vector or Flag-Hsp70 were exposed to heat shock at 45°C for 15 min and recovered at 37°C for the indicated times. The overexpression of Hsp70 in the HSF1-/- cells facilitated the degradation of the Arc/Arg3.1 protein by about 50% (Fig. 2B and 2C), suggesting that Hsp70 plays a role in the degradation of the Arc/Arg3.1 protein. To determine whether Arc/Arg3.1 is the binding substrate of Hsp70, we examined the interaction between Hsp70 and Arc/Arg3.1. HEK293T cell lysates transfected with GFP-Hsp70 and Flag-control or Flag-Arc were immunoprecipitated with an anti-Flag antibody, and we found that Arc/Arg3.1 is associated with GFP-Hsp70 (Fig. 2D). The endogenous interaction between Arc/Arg3.1 and Hsp70 was confirmed in the radiation-induced mouse fibrosarcoma cell line, RIF-1, in which Hsp70 is highly expressed in response to heat shock (Fig. S1A). These interactions show the possibility that heat shock-induced Arc/Arg3.1 degradation occurrs through Hsp70 chaperone-assisted UPS. Since heat shock caused only slight expression of Hsp40 and Hsp27 in HSF1-/- cells as shown in Fig. 2A, we investigated the potential link between Arc/Arg3.1 and either Hsp40 or Hsp27. Immunoprecipitation experiments with Flag-Arc revealed that Arc/Arg3.1 primarily interacted with Hsp70 (Fig. 2E). Therefore, we were able to rule out the hypothesis that chaperone-assisted UPS mediated by Hsp40 or Hsp27 is responsible for Arc/Arg3.1 degradation.

**Fig. 2.**
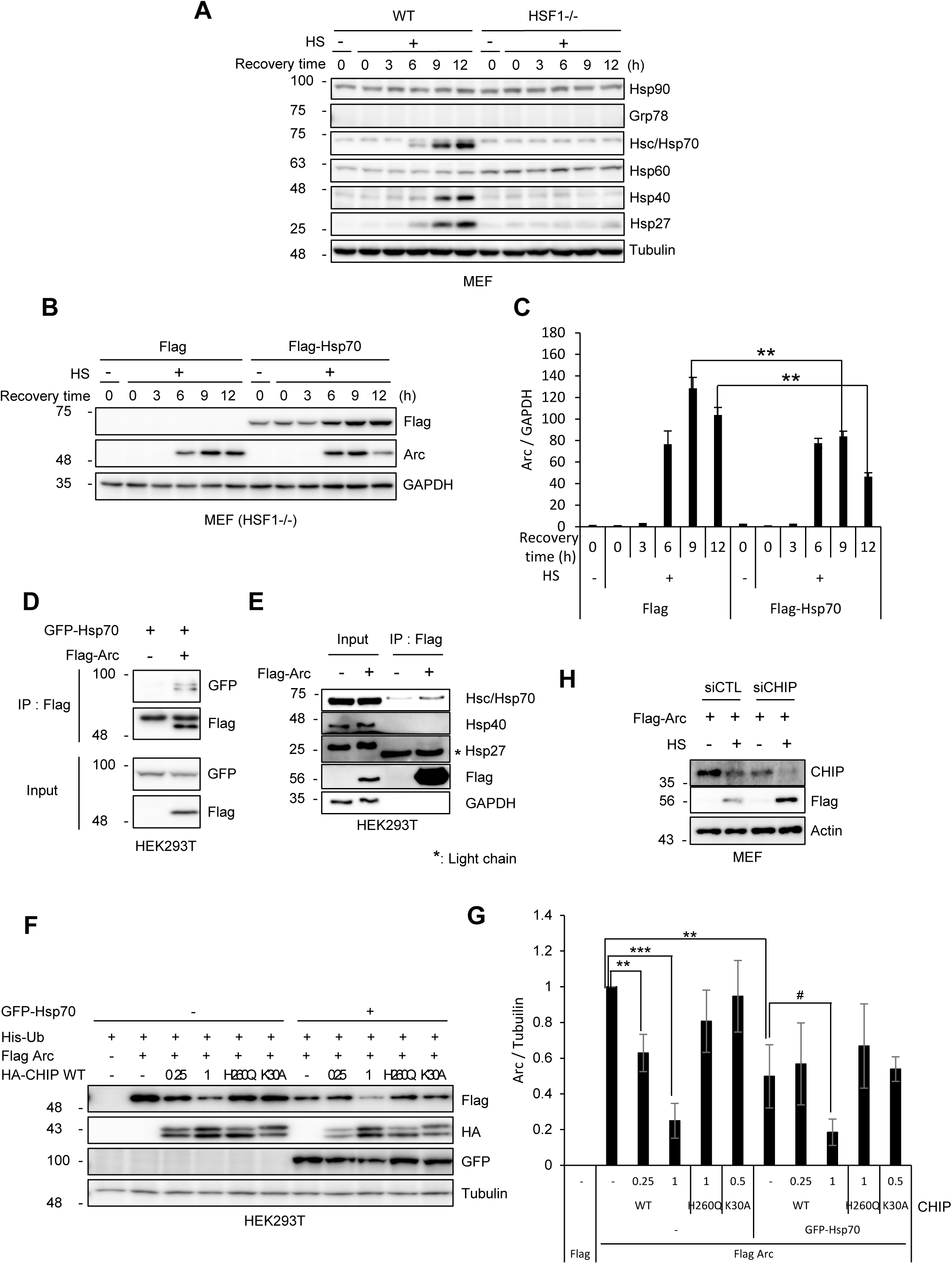
Hsp70 interacts with Arc/Arg3.1 and facilitates the degradation of Arc/Arg3.1. (A) HSF1-/- cells exhibit reduced expression of heat shock inducible Hsps. WT and HSF1-/- MEF cells were exposed to heat shock at 45 °C for 15 min and allowed to recover at 37 °C for the indicated times. Hsp90, Grp78, Hsc/Hsp70, Hsp60, Hsp40, and Hsp27 were detected by Western blot analysis with their specific antibodies. (B, C) Hsp70 participates in UPS-dependent Arc/Arg3.1 degradation. HSF1-/- MEF cells overexpressing Flag-Hsp70 were exposed to heat shock at 45 °C for 15 min and allowed to recover at 37 °C for the indicated times. Flag-Hsp70 and Arc/Arg3.1 were detected by Western blot analysis using their specific antibodies. Representative Western results were selected from triplicated experiments. Quantified results of triplicated Western images are shown in (C). (D) Hsp70 interacts with Arc/Arg3.1. HEK293T cells were transfected with GFP-Hsp70 and Flag empty vector or Flag-Arc, and the cell lysates were immunoprecipitated using an anti-Flag antibody. The immune complex was analyzed by Western analysis using anti-GFP and anti-Flag antibodies. (E) Arc/Arg3.1 primarly binds to Hsc/Hsp70. Following transfection of HEK293T cells with either Flag empty vector or Flag-Arc, the whole cell lysates were immunoprecipitated using an anti-Flag antibody. Western analysis using anti-Hsc/Hsp70, anti-Hsp40, and anti-Hsp27 antibodies was used to examine the immunological complex. * was an antibody’s light chain. (F, G) CHIP induces Arc/Arg3.1 degradation by interacting with Hsp70. HEK293T cells overexpressing Flag-Arc, GFP-Hsp70, and His-ubiquitin were co-transfected with HA-CHIP WT or mutants. Cells were analyzed by Western blot with anti-Flag, anti-HA, and anti-GFP antibodies. Representative Western results were selected from triplicated experiments. Quantified results of triplicated Western images are shown in (G). Data are presented as the mean ± S.D. of triplicated experiments (t-test; # < 0.01, **P < 0.001, *** < 0.0001). (H) Arc/Arg3.1 proteins are stabilized by CHIP knockdown. After being depleted of either control or CHIP, MEF cells were exposed to heat shock at 45°C for 15 min and then allowed to recover at 37°C for 9 h.

To further investigate whether Hsp70 associated E3 ligase CHIP induces the degradation of Arc/Arg3.1, we examined the degradation of Arc/Arg3.1 using various functional mutants of CHIP. HEK293T cells overexpressing Flag-Arc, GFP-Hsp70 and His-ubiquitin, were co-transfected with CHIP WT, H260Q mutant (enzymatic activity-deficient mutant), or K30A mutant (Hsp70 interaction deficient mutant), and the Arc/Arg3.1 protein levels were assessed. As shown in Fig. 2F, Arc/Arg3.1 was more readily degraded by increasing the CHIP WT expression. In contrast, Arc/Arg3.1 was not degraded in cells overexpressing either CHIP mutant. This result shows that CHIP plays a role in degrading Arc/Arg3.1 by interacting with the chaperones. To confirm the effect of Hsp70 on Arc/Arg3.1 degradation by CHIP, GFP-Hsp70 was overexpressed in the same conditions as the above experiment. Arc/Arg3.1 was more readily degraded in cells overexpressing Hsp70 by CHIP WT than by the CHIP mutants (Fig. 2F and 2G). The results indicate that the CHIP-Hsp70 complex is required to induce Arc/Arg3.1 degradation. Consequently, CHIP depletion resulted in the stabilization of Arc/Arg3.1 proteins (Fig. 2H). Additionally, we confirmed that Arc/Arg3.1 degradation mediated by the CHIP-Hsp70 complex is proteasome-dependent, as MG132 treatment effectively prevented the degradation of Arc/Arg3.1 in the presence of overexpressed CHIP and Hsp70 (Fig. S1B).

### 2.3. Disulfide bonds contribute to the aggregation and oligomeric property of Arc/Arg3.1

Numerous efforts have been made to elucidate the full-length structure of Arc/Arg3.1. However, significant challenges remain in achieving crystallization due to the flexible regions within the Arc/Arg3.1 protein and its inherent tendency toward aggregation and oligomerization, particularly in the N-terminal region (39, 43, 44). Purified recombinant Arc/Arg3.1 protein is easily aggregated into turbid solutions at high protein concentrations (>1 mg/mL). However, the Arc/Arg3.1 band appeared at the monomer position (55 kDa) under reducing SDS-PAGE conditions with β-mercaptoethanol (β-ME). Recently, it was confirmed that mammalian brain contains Arc/Arg3.1 oligomers, and *in situ* crosslinked low-order oligomer band was intensified in non-reducing SDS-PAGE (45), suggesting that Arc/Arg3.1’s quaternary structures involve the creation of disulfide bonds.

In order to determine whether Arc/Arg3.1 generates oligomers via disulfide bond, we examined the molecular weight of Arc/Arg3.1 under non-reducing SDS-PAGE without β-ME. Arc/Arg3.1 bands were observed at the dimer, trimer, and high-order multimers, with aggregates in the stacking gel under non-reducing conditions, unlike the monomers under the reducing condition (Fig. 3A, left panel). By using anti-Arc antibody for immunoblotting, we were able to demonstrate that these bands were, in fact, Arc/Arg3.1 (Fig. 3A, right panel). The oligomers and aggregates of the recombinant Arc/Arg3.1 proteins were readily reduced to monomers by reducing agent DTT in concentration-dependent manner (Fig. S2A). The size of the recombinant Arc/Arg3.1 proteins was further confirmed using Dynamic Light Scattering (DLS), revealing an Arc/Arg3.1 oligomer with a size of 30–40 nm, consistent with previous findings (44, 46). Endogenous Arc/Arg3.1, induced in HEK293T cells exposed to heat shock at 45°C for 15 min and recovered at 37°C for the indicated times, were fractionated into soluble (S) and insoluble (I) fractions and separated on SDS-PAGE with and without β-ME. The soluble fraction of cells shows the monomer having an intra-disulfide bond, presented as an oxidized monomer, which moves faster than a reduced monomer in non-reduced SDS-PAGE, whereas the insoluble fraction is mainly oligomers (Fig. 3C). The results indicate that disulfide crosslinkings are key factor to generate Arc/Arg3.1 oligomer both in recombinant and cellular proteins.

**Fig. 3.**
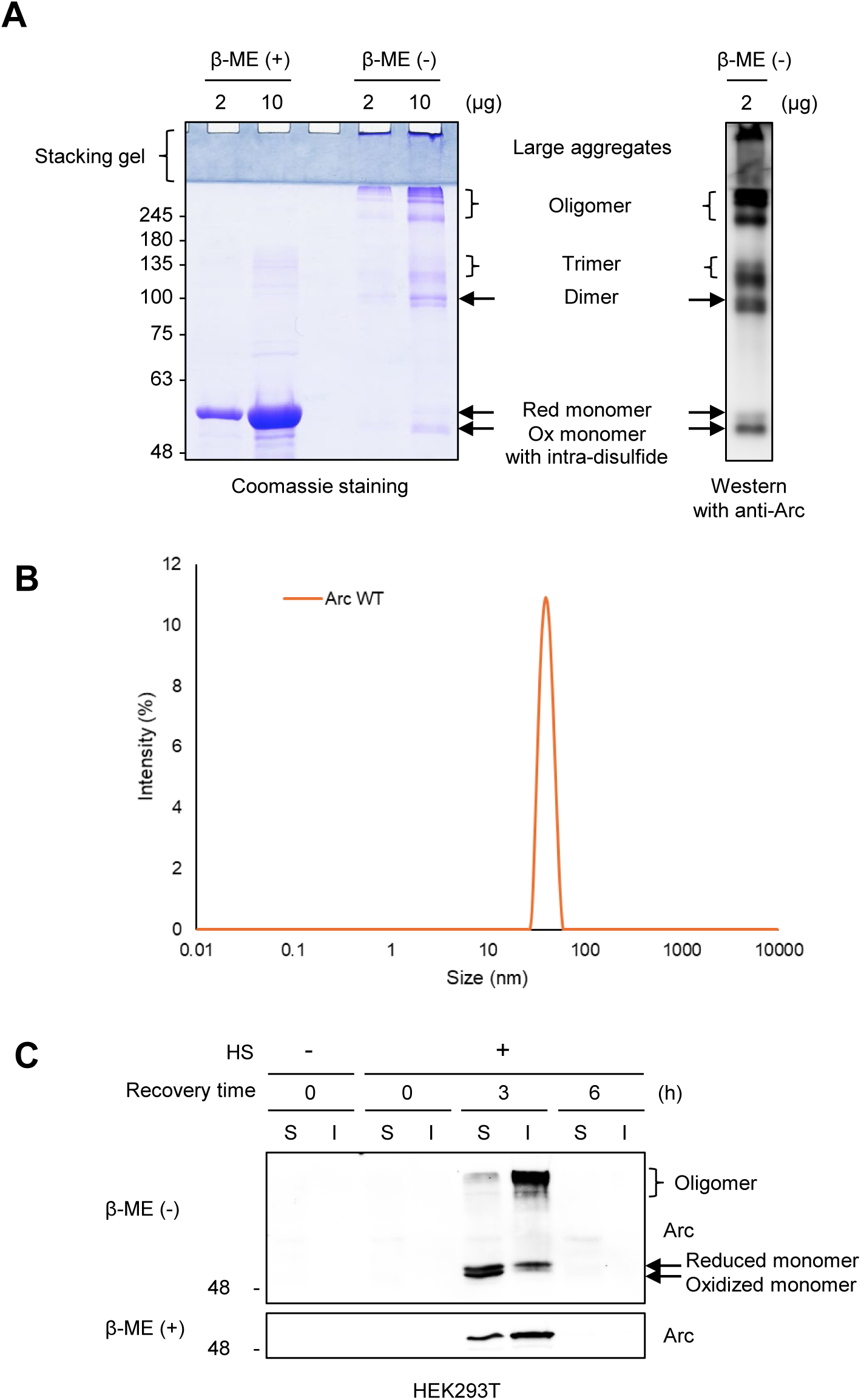
Disulfide bond formation contributes to the aggregation and oligomeric property of Arc/Arg3.1. (A) Purified hArc/Arg3.1 proteins were separated by SDS-PAGE under reducing (+) β-ME) and non-reducing (-β-ME) conditions, and stained with coomassie blue staining (left panel) and detected by Western blot analysis using an anti-Arc antibody (right panel). (B) The size of recombinant Arc/Arg3.1 protein was analyzed by DLS. (C) HEK293T cells were exposed to heat shock at 45 °C for 15 min and allowed to recover at 37 °C for the indicated times, then fractionated into soluble (S) and insoluble (I) fractions. The proteins in each fraction were separated by SDS-PAGE under non-reducing (-β-ME) and reducing (+ β-ME) conditions and detected by Western analysis using anti-Arc antibodies.

### 2.4. Arc/Arg3.1 forms disulfide bonds between Cys34 and Cys159

Human Arc/Arg3.1 has five cysteine residues, all located in the N-terminal half (aa 1-206), four Cys residues in the matrix-like domain (MA, aa 17 ∼ 154), and one Cys in the linker (Fig. 4A). To identify the disulfide bonds involved in Arc/Arg3.1 oligomerization, we generated three cysteine to alanine mutants (C34A, C94/98A, C159A) and examined the oligomerization under non-reducing conditions. All cysteine mutants showed reduced large oligomer formations compared to the WT, and oligomeric forms are readily reduced to monomers, including intra-disulfide linkage with low concentrations of DTT (Fig. 4B and 4C). Predominantly, both the C34A and C159A mutants exist in the monomer state, while the C94/98A mutant primarily forms dimers and oligomers, suggesting that Cys34 and Cys159 are involved in disulfide bond formation and play critical roles in the initial oligomerization. To identify the intact disulfide bonds of the Arc/Arg3.1 dimer and aggregates, we employed the methodology to analyze the disulfide linkage by peptide sequencing with nanoUPLC-ESI-q-TOF mass spectrometry combining disulfide searching algorithm DBond(38). Cys34-Cys159 disulfide crosslinkings were identified in aggregate band of the WT Arc/Arg3.1 (band 1 in Fig. 4B) and the dimer band of the C94/98A mutant (band 2 in Fig. 4B). The intact MS/MS spectrum of the Cys34-Cys159 disulfide peptides was achieved by digesting the long disulfide peptides with trypsin combined with chymotrypsin to a shorter one (^34^C*RAEMLEHVR^43^-^159^C*HEADGYDY^167^ (m/z = 1005.4292 Da, z = 3)) to raise the ionization efficiency (Fig. 4D). Since the next residue of Cys34 is arginine, the digested peptide by trypsin is too short to be searched for by the analysis program. Therefore, by manually searching with the unknown modification search tool, we obtained the peak with ^34^CR^35^ bound to C159 (^34^C*R^35^-^159^C*HEADGYDY^167^) (Fig. S4). In addition, various intra-disulfide linkages among Cys94, Cys96, and Cys98 (^93^ACLCRCQETIANLER^107^) were identified in the aggregated band 1 of the WT Arc/Arg3.1 (Table S1). The results indicate that disulfide bond formations play an essential role in inducing the structural changes of Arc/Arg3.1 to the aggregated form.

**Fig. 4.**
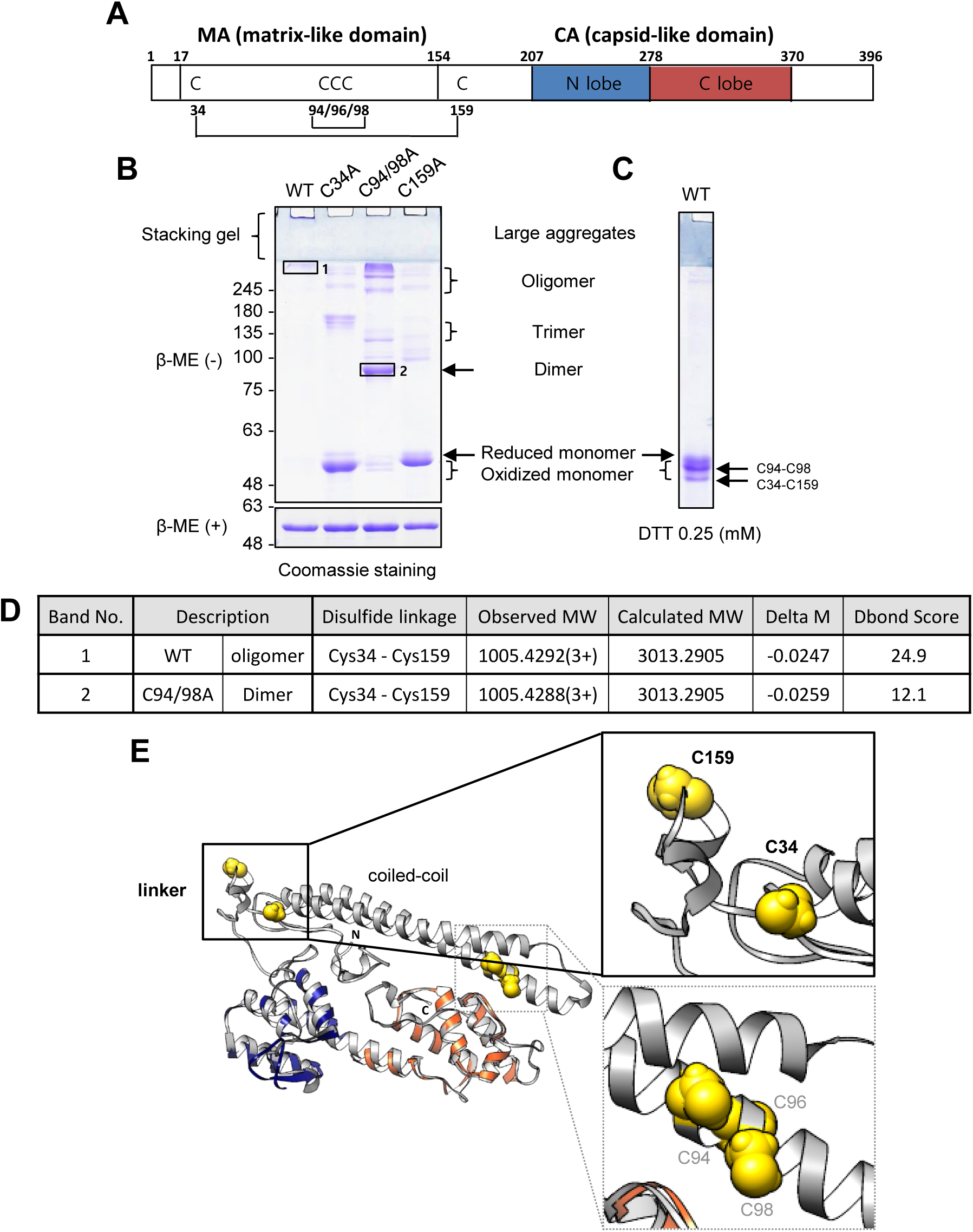
Arc/Arg3.1 forms disulfide bond between Cys34 and Cys159. (A) Schematic of hArc/Arg3.1 domains and location of cysteine (C) residues. (B) Purified hArc/Arg3.1 WT and cysteine mutant (C34A, C94/98A, and C159A) proteins pre-treated with 20 mM NEM at 37 °C for 10 min were separated by SDS-PAGE gel under non-reducing (-β-ME) and reducing (+ β-ME) conditions and stained with coomassie blue. (C) Purified hArc/Arg3.1 protein treated with 0.25 mM DTT at 37 °C for 30 min were separated by SDS-PAGE under non-reducing conditions. (D) Disulfide bonds of labeled protein bands in (B) were analyzed by peptide sequencing with nanoUPLC-ESI-q-TOF tandem mass spectrometer combined with the DBond searching algorithm. (E) The full-length hArc/Arg3.1 model was obtained from the Robetta protein structure service (http://new.robetta.org). Superposition of modeled structure of full-length Arc/Arg3.1 (from Robetta: comparative modeling; grey) with Arc/Arg3.1 N-lobe (PDB: 4X3H; navy) and C-lobe (PDB: 4X3X; orange).

To investigate how disulfide formation alters the 3D structural changes, we examined the structure of the full-length Arc/Arg3.1 by modeling the structure using a structure prediction program (Robetta, http://new.robetta.org); owing to the aggregation properties of the protein, the X-ray crystal structure of the full-length Arc/Arg3.1 is curruently unavailable. The modeled structure of full-length Arc/Arg3.1 is very similar to the recently presented hybrid model (combining small-angle X-ray scattering (SAXS) data, the crystal structure of the CA domain and the homology model of the N-terminal region)(39) (Fig. 4E). The structure suggests that Cys34 and Cys159 are adjacent with an appropriate distance to form an intra-disulfide bond. Cys94, Cys96, and Cys98 (CXCXC motif) can readily form intra-disulfide bonds with adjacent cysteine residues. The Arc/Arg3.1 WT protein appears as oligomers and aggregates under non-reducing conditions, yet the inter-disulfide crosslinkings are readily reduced into three monomer bands by mild reducing conditions (<0.25 mM DTT). Whereas C159A mutants mostly showed a reduced monomer band, Arc/Arg3.1 WT monomer bands include both intra-disulfide bonds (Cys94-Cys98 and Cys34-Cys159) (Fig. S2B). Comparison to the monomer band position of the cysteine mutants, the band just below the reduced monomer is assumed to be the Cys94-Cys98 intra-disulfide bond, and the bottom band is the Cys34-Cys159 intra-disulfide bond (Fig. 4B, C). The location of the cysteine residues in the 3D structure is well agreed with the intra- and inter-disulfide bonds on the non-reducing gel.

### 2.5. Cys159 is a reactive cysteine in Arc/Arg3.1 and is readily oxidized by oxidative stress

In order to investigate which cysteine residues play crucial roles in the disulfide formation, we examined the reactivities of the cysteine residues in the recombinant Arc/Arg3.1 proteins, employing chemical probe NPSB-B, a novel biotin-labeling probe that specifically reacts with redox-sensitive Cys-SH residues (47). Recombinant Arc/Arg3.1 WT proteins purified with reducing agent DTT were incubated with various concentrations of H_2_O_2_ for 1 h and then labeled with NPSB-B for 2 h. WT Arc/Arg3.1 was readily labeled with NPSB-B, and then the labeling was decreased upon H_2_O_2_ treatment in a dose-dependent manner, indicating that WT Arc/Arg3.1 has redox sensitive cysteine residue(s) (Fig. 5A). The C34A and C94/98A mutants were labeled with NPSB-B as well as WT, while the C159A mutant labeling was significantly decreased (Fig. 5B and 5C). The result indicates that Cys159 is the most redox sensitive cysteine residue. The results were confirmed by incubation with various concentrations of H_2_O_2_. The NPSB-B labelings of Arc/Arg3.1 WT, C34A, and C94/98A were decreased in an H_2_O_2_ concentration-dependent manner. However, the C159A mutant could not be labeled with NPSB-B (Fig. 5D and 5E). This indicates that Cys159 is the redox sensitive cysteine residue, and the reactivity of Cys159 in WT, C34A, and C94/98A mutants is decreased by oxidation following H_2_O_2_ treatment. We subsequently used an immunofluorescence (IF) assay to compare the staining patterns of the Arc/Arg3.1 WT and the C159A mutant in order to better comprehend their oligomeric properties in cellular system. Whereas the Flag-Arc C159A mutant showed a scattered and dispersed staining over the entire cell, the Flag-Arc WT revealed protein aggregates (Fig. S3C). The results suggest that the Cys159 residue is a nucleophile with reactivity to form disulfide crosslinking with Cys34, contributing to Arc/Arg3.1 self-assembly.

**Fig. 5.**
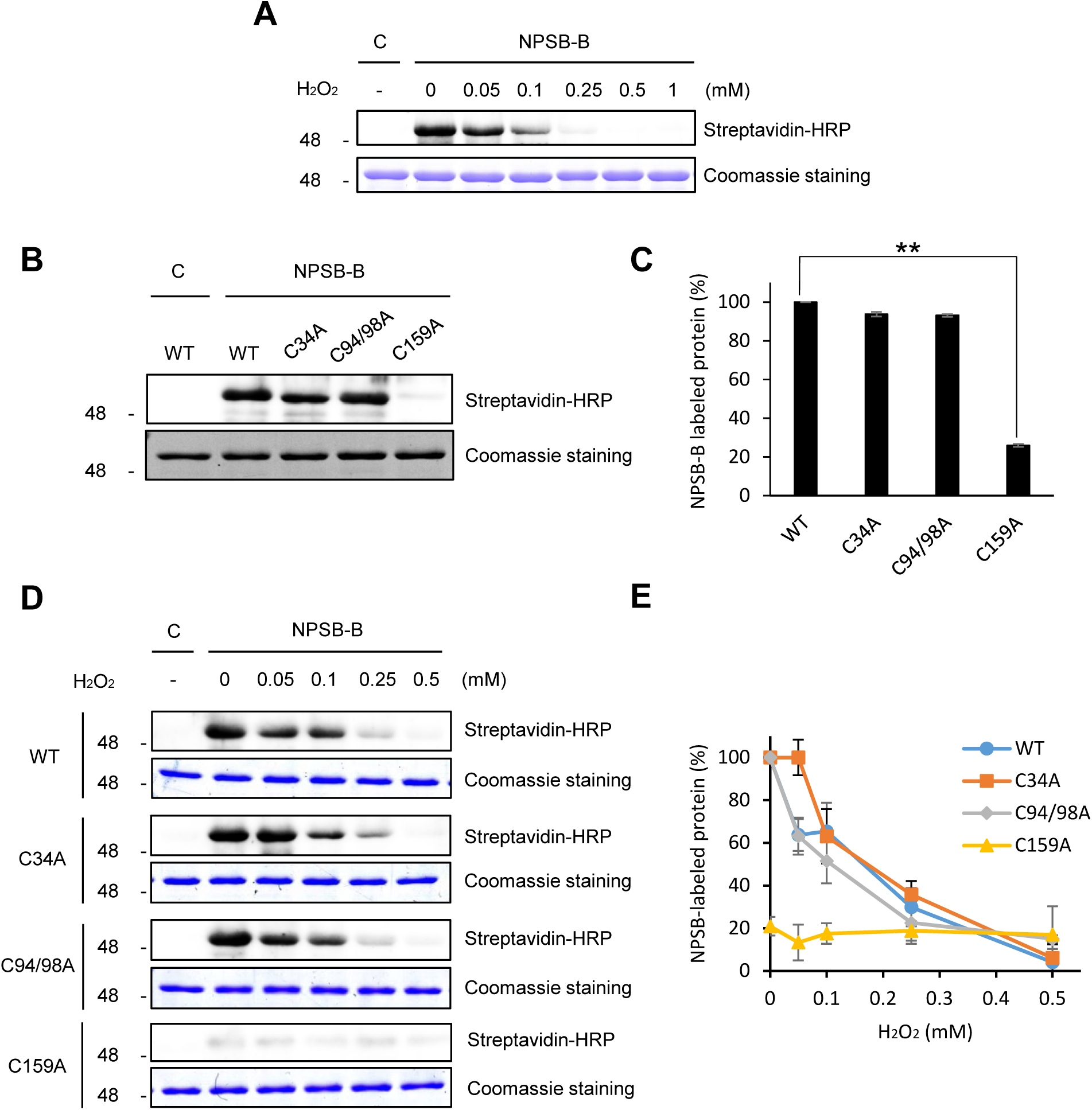
Cys159 is a reactive cysteine residue of Arc/Arg3.1 and is readily oxidized by oxidative stress. (A) Purified hArc/Arg3.1 protein prepared under reducing conditions was treated with indicated concentrations of H_2_O_2_ at 37 °C for 1 h and then labeled with 1 mM NPSB-B at RT for 2 h. NPSB-B labeled samples in gel sample buffer containing 20 mM NEM were detected by streptavidin-HRP (upper panel), and loaded protein amounts were detected using coomassie blue staining (lower panel). (B, C) Arc/Arg3.1 WT and cysteine mutants (C34A, C94/98A, and C159A) were labeled with 1 mM NPSB-B at RT for 2 h. NPSB-B labeled samples were detected by streptavidin-HRP (upper panel), and loaded protein amounts were detected using coomassie blue staining (lower panel). Amounts of NPSB-B labeled proteins were quantified (C). Data are presented as the mean ± S.D. of triplicated experiments (t-test; **P < 0.01). (D, E) Arc/Arg3.1 WT and cysteine mutants (C34A, C94/98A, and C159A) were treated with indicated concentrations of H_2_O_2_ at 37 °C for 1 h and then labeled with 1 mM NPSB-B at RT for 2 h. NPSB-B labeled samples were detected by streptavidin-HRP (upper panel) and coomassie staining (lower panel) (D). Labeled amounts were quantified following triplicated experiments (E).

Cysteine residues are characterized by either high or low conservation (48). To understand the evolutionary meaning of each cysteine residue in the Arc/Arg3.1 protein, the cysteine conservations were confirmed through sequence alignment. Five cysteines in the human Arc/Arg3.1 are almost all conserved in mammals (Table S2). Cys94 and Cys98, which are conserved throughout the vertebrate lineage, are present in the ^94^CLCRC^98^ motif. Reactive Cys159 in the flexible linker appears to be conserved in mammals, yet the structural prediction program confirms that analogous cysteine is present in some species of birds and reptiles and that they are all adjacent to Cys34 in the tertiary structure. It is regarded that functional cysteines are conserved through evolution (48). Cysteine in the C^94^LC^96^RC^98^ motif, known for its membrane lipid binding via palmitoylation (49), has been conserved since Arc/Arg3.1 was inserted early in the vertebrate lineage. Reactive Cys159 might be acquired as it evolved to higher vertebrates, which appears to be conserved due to the requirement of a regulatory function. The evolutionary acquisition of redox-sensitive cysteine and disulfide bond implies the potential of redox regulation as a main protein regulation system.

### 2.6. Disulfide formation via Cys159 is a prerequisite for the degradation of Arc/Arg3.1 via CHIP dependent ubiquitination

The structural changes caused by disulfide bonds can affect protein stability (34, 50). Moreover, reactive oxygen species (ROS) levels increase in response to various cellular stressors that lead to Arc/Arg3.1 induction and degradation (10). We hypothesized that disulfide crosslinking may regulate the oligomeric state of Arc, thereby influencing its proteasomal degradation, as the oxidation of cysteine residues is a common post-translational modification in response to elevated ROS levels (51). To investigate whether disulfide formation affects the stability of Arc/Arg3.1, we examined the degradation kinetics of Arc/Arg3.1 WT and C159A mutant. HEK293T cells transiently transfected with Flag-Arc WT or C159A mutant were treated with cycloheximide (CHX), an inhibitor of mRNA translation, for indicated times. We found that the half-life of the WT Arc/Arg3.1 was 6 h, while the C159A mutant was not degraded in 6 h (< 10%) (Fig. 6A and 6B). This indicates that oligomerized or aggregated Arc/Arg3.1 via Cys159-Cys34 crosslinking are required for the degradation of Arc/Arg3.1.

**Fig. 6.**
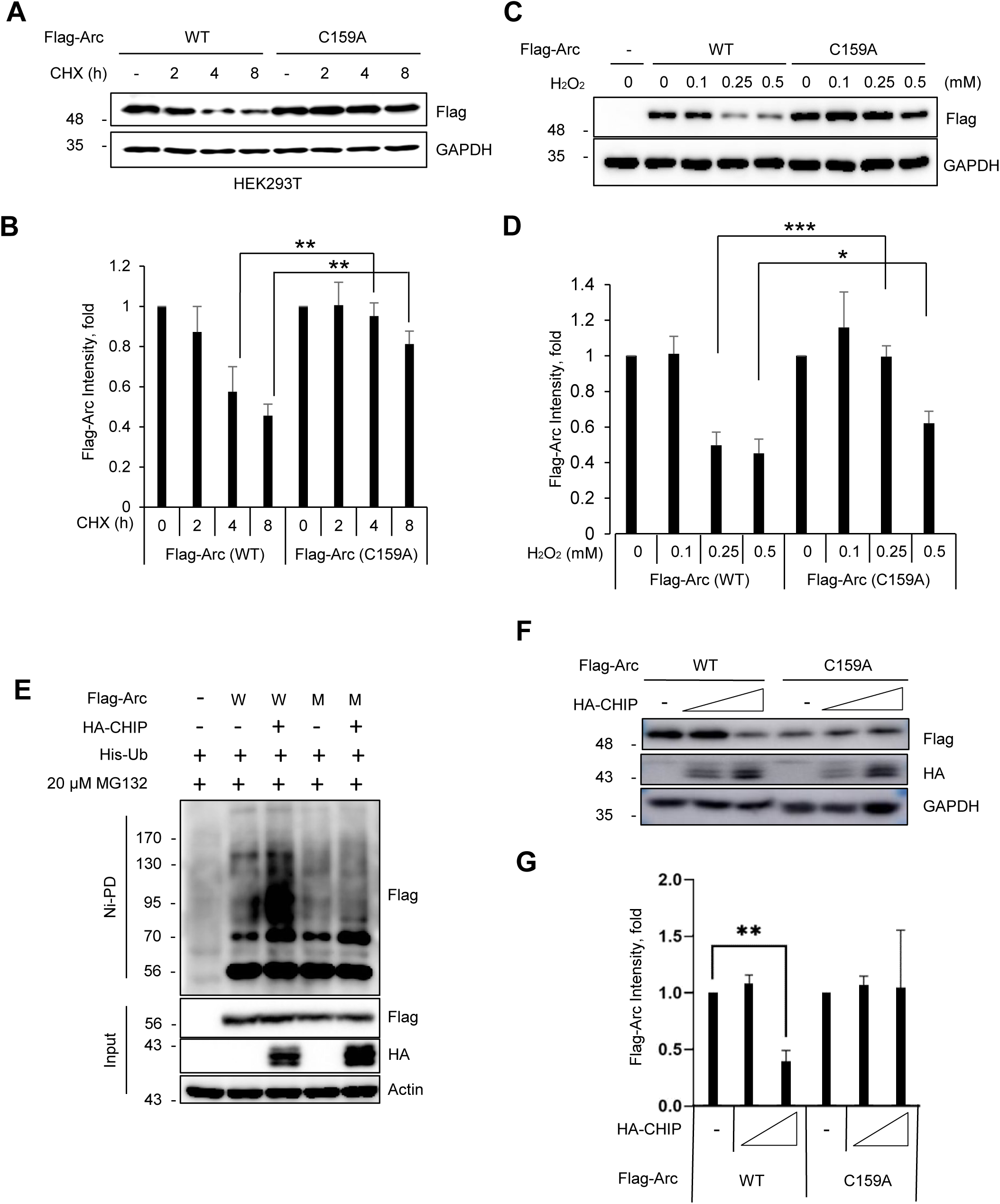
Disulfide formation is required for ubiquitination and degradation of Arc/Arg3.1. (A, B) The lifetime of the Arc/Arg3.1 C159A mutant is much longer than that of WT. HEK293T cells overexpressing Flag-Arc WT or C159A mutant were treated with 10 μg/mL cycloheximide (CHX) for the indicated times. The Arc/Arg3.1 protein degradation was analyzed by Western blotting using an anti-Flag antibody. GAPDH was used as a loading control. Representative Western results were selected from triplicated experiments. Quantified results of quadruplicated Western images are shown in (B). Data are presented as the mean ± S.D. of triplicated experiments (t-test; **P < 0.01). (C, D) HEK293T cells overexpressing Flag-Arc WT or C159A mutant were treated with the indicated concentrations of H_2_O_2_ at 37 °C for 1 h, followed by treatment with 10 μg/mL cycloheximide (CHX) for 4 h. Cell lysates were separated by SDS-PAGE and detected by Western blot analysis using anti-Flag antibody. Quantified results of quadruplicated Western images are shown in (D). Data are presented as the mean ± S.D. of triplicated experiments (t-test; **P < 0.01). (E) For 4 h, 20 μM MG132 was applied to HEK293T cells that were overexpressing His-Ub, HA-CHIP, and either Flag-Arc WT or C159A in order to prevent UPS-mediated degradation. Following Ni-NTA agarose precipitation of cell lysates, the ubiquitinated Arc proteins within the immunological complex were identified via Western blot analysis employing an anti-Flag antibody. (F, G) HEK293T cells transfected with two concentrations of HA-CHIP (0.25 μg and 0.5 μg) and Flag-Arc WT or C159A were incubated for 24 h, and degradation of the different Flag-Arc proteins was assessed by Western blot analysis using anti-Flag and anti-HA antibodies. GAPDH was used as a loading control. Quantified results of quadruplicated Western images are shown in (G). Data are presented as the mean ± S.D. of triplicated experiments (t-test; **P < 0.01).

To investigate whether degradation of WT Arc is redox-regulated, we examined the stability of Arc/Arg3.1 WT and C159A under oxidative stress. Arc/Arg3.1 WT was degraded but C159A mutant remained more stable even under oxidative stress (Fig. S3A, B). Subsequently, we investigated whether overexpressed Arc/Arg3.1 WT and C159A mutant are recovered when cells are incubated with CHX for 4 h after treatment with various concentrations of H_2_O_2_ for 1 h. While WT Arc/Arg3.1 was degraded in 0.25 mM H_2_O_2_, the C159A mutant remained stable (Fig. 6C and 6D). The results suggest that degradation of the WT Arc/Arg3.1 is regulated by the redox-sensitive oligomerization.

To determine whether Arc/Arg3.1 ubiquitination can be differentiated between WT and C159A mutant, we evaluated the ubiquitination of Arc/Arg3.1 caused by CHIP, which has been identified as an E3 ligase for Arc/Arg3.1 (16). We overexpressed His-Ub, HA-CHIP, and either Flag-Arc WT or C159A mutant in the presence of MG132 for 4 hours. Ubiquitin conjugates were then isolated using nickel agarose beads. The results showed that WT was successfully ubiquitinated by CHIP, while the C159A mutant was not (Fig. 6E). Moreover, we examined whether the Arc/Arg3.1 WT and C159A mutant undergo distinct degradation by CHIP. CHIP led to a dose-dependent decrease in WT Arc/Arg3.1 expression, yet the C159A mutant expression remained unaffected (Fig. 6F and 6G). The results show that WT Arc/Arg3.1 is readily ubiquitinated by E3 ligase and degraded by the UPS. However, the C159A mutant, which does not form oligomeric and aggregated forms via disulfide bond, remains stable and is not degraded by the UPS. These data suggest that termination of Arc/Arg3.1 function by rapid degradation is tightly regulated by the production of oligomeric and aggregated Arc/Arg3.1 via Cys159-Cys34 disulfide formation.

### 2.7. Cys159-Cys34 intra-disulfide bond in Arc/Arg3.1 plays a crucial role in regulating its structure

To investigate whether Cys159-Cys34 intra-disulfide bond affects the Arc/Arg3.1 structure, we examined the protein structural differences by employing hydrogen-deuterium exchange mass spectrometry (HDX-MS) (52). HDX-MS was performed as described in the methodology and is summarized in Table S3. The H/D exchange rates for the WT and C159A mutant were compared for each pepsin-digested peptide (Fig. 7A, Table S4). Combined stitching H/D exchange rates of pepsin-digested peptides showed the diagram of the whole protein (Fig. 7A). The peptide MS coverage of the Arc/Arg3.1 WT was 71%, while the MS coverage of the soluble proteins is commonly 100% (34, 50) (Table S3). The WT Arc/Arg3.1 is thought to have low coverage because it cannot be sufficiently digested by pepsin. Conversely, the C159A mutant is less aggregated, with a coverage of 84%. Although the sequence coverage for the Arc/Arg3.1 protein is low and the interpretation of the results is limited, these results show that the significant H/D exchange rate differences between the WT and C159A mutant appeared in critical protein regions. We mapped the Arc/Arg3.1 sequence onto the full structure modeled by Robbeta (Fig. 7B). For accurate analysis of the N-lobe of the CTD, we used the rat Arc/Arg3.1 crystal structure (53) to display the results of the H/D exchange rates (Fig. 7C and D).

**Fig. 7.**
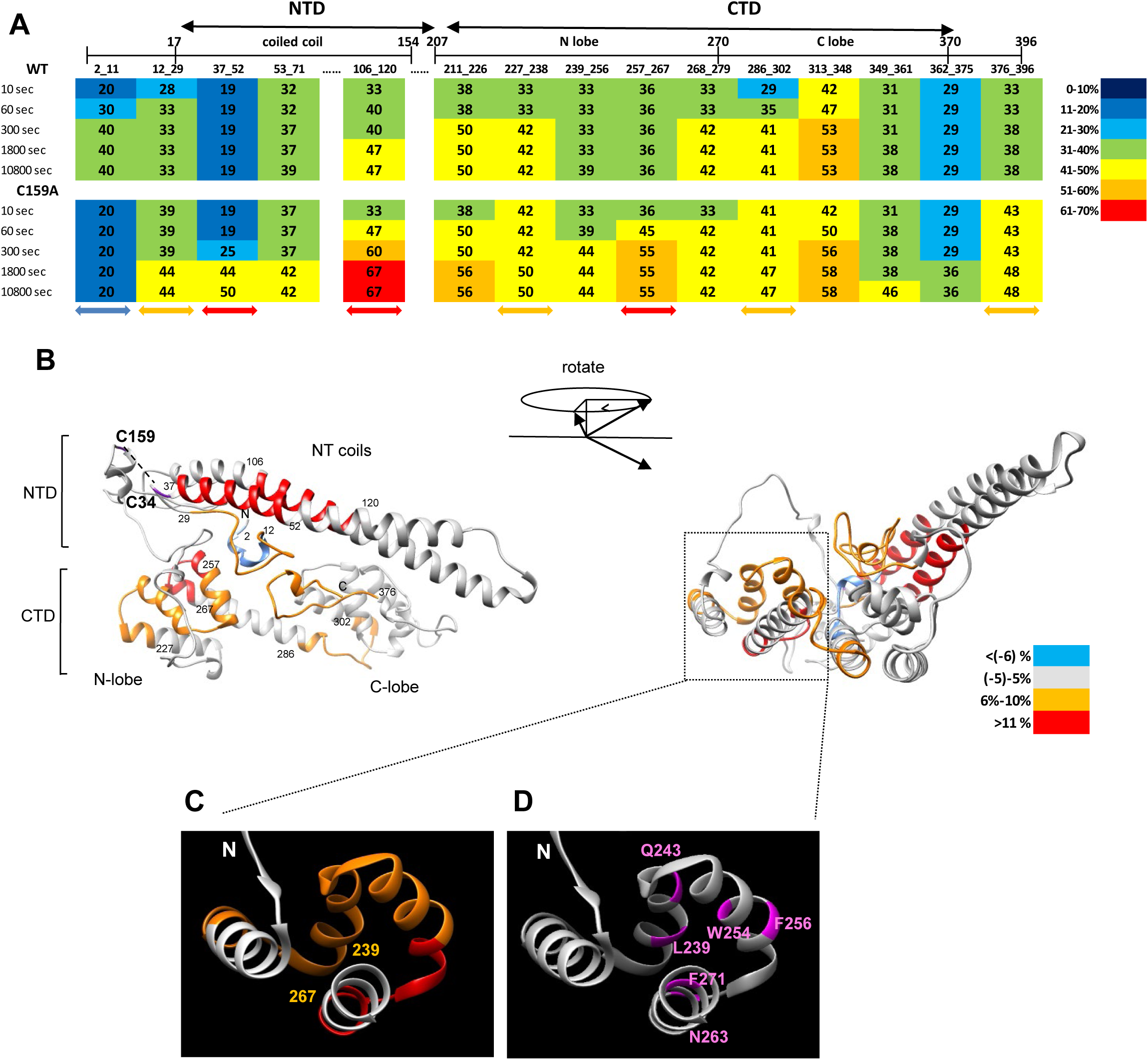
Identification of structural differences between hArc/Arg3.1 WT and C159A mutant employing hydrogen/deuterium exchange mass spectrometry (HDX-MS). (A-B) Recombinant WT and C159 mutant Arc/Arg3.1 proteins were incubated with D_2_O exchange buffer at 25 °C for various times up to 30 min and analyzed using nanoAcquity^TM^/ESI/MS. (A) The deuterium exchange rate (%) of each peptide in WT and C159A mutant Arc/Arg3.1 was presented depending on the D_2_O incubation time. Colored arrows mark the significant differences between the WT and C159A mutant. (Differential deuterium exchange rate of C159A compared to WT; blue: ≥ -6%, orange: 6%-10%, and red: 11% or more). The corresponding deuterium exchange levels for each peptide are given in percentages on the right. (B) Overlay differential HDX data onto the hArc/Arg3.1 structure modeled from Robetta. This overlay compares the HDX of WT Arc/Arg3.1 with that of the C159A mutant. (C) Differential HDX data overlay on the crystal structure of a specific site, rArc /Arg3.1 N-lobe (PDB: 4X3H). (D) Six residues in the rArc/Arg3.1 N-lobe (PDB: 4X3H) caused significant conformational changes by GluN2A (NMDA receptor).

The C159A mutant exhibited a higher deuterium exchange rate than the WT in most areas, indicating that the C159A mutant is structurally more open and exposed to solvent. The H/D exchange levels of a peptide containing coiled-coil inlet (37–52), the middle of coiled-coil (106–120), and some regions in the N-lobe (239–267) were significantly increased in the C159A mutant (Fig. 7B). The deuterium exchange rate of the peptide (106–120), which harbors the oligomerization motif (113–119) critical for Arc/Arg3.1 self-association (37), was significantly increased in the C159A mutant. This observation suggests that the structural stability of Arc/Arg3.1 may be compromised in this mutant. In the overall structure, the exchange level increased near the interface between the N-terminal domain (NTD, 1-140) and the C-terminal domain (CTD, 208-396) in the C159A mutant. Thus, the loss of Cys34-Cys159 crosslinking in the C159A mutant appears to attenuate the interaction between NTD and CTD. Overall, the disulfide bonds play an essential role in proper folding of Arc/Arg3.1 and domain orientation during oligomerization. These dynamic structural analyses indicated that the C159A mutant undergoes discernible changes to a more soluble structure, causing it to lose its degradation properties of as a key regulation system.

### 2.8. The stable Arc/Arg3.1(C159A) mutant abolished the negative effects on Hsp70 expression by losing the interaction with HSF1

Previous study has shown that Arc/Arg3.1 binds to HSF1 and prevents it from binding to HSEs, thereby reducing Hsp70 induction in non-neuronal system (10). We observed distinct degradation machinery, originated from oligomeric properties between WT Arc/Arg3.1 and C159A mutant so far, the functional difference remains unclear. To investigate the effect on Hsp70 expression, cells transfected with Arc/Arg3.1 WT or the C159A mutant were exposed to heat shock and left to recover for the indicated time and Hsp70 levels were assessed. Arc/Arg3.1 WT overexpression led to decreased Hsp70 expression compared to the control, whereas the C159A mutant did not show negative effects on Hsp70 expression (Fig. 8A and 8B). Subsequently, the subcellular localization of the Flag-Arc WT and C159A mutant was investigated since activated HSF1 is known to be localized in the nucleus. HEK293T cells overexpressing Flag-Arc WT or Flag-Arc C159A were exposed to heat shock and fractionated into cytosolic and nuclear fractions. Both the Arc WT and C159A mutants showed nuclear translocation in response to heat shock (Fig. 8C). Thus, further analysis examined the interaction between the Arc WT and C159A mutant and HSF1. As shown in Fig. 8D, only Flag-Arc WT and not the Flag-Arc C159A mutant interacted with HSF1, indicating that the stable C159A mutant abolished the negative effects on HSF1 transcriptional activity by losing the interaction with HSF1. Hence, our study revealed a crucial double-negative feedback loop involving Arc, HSF1, and HSP70, which is pivotal in governing the cellular response to stress, such as oxidative stress or heat shock.

**Fig. 8.**
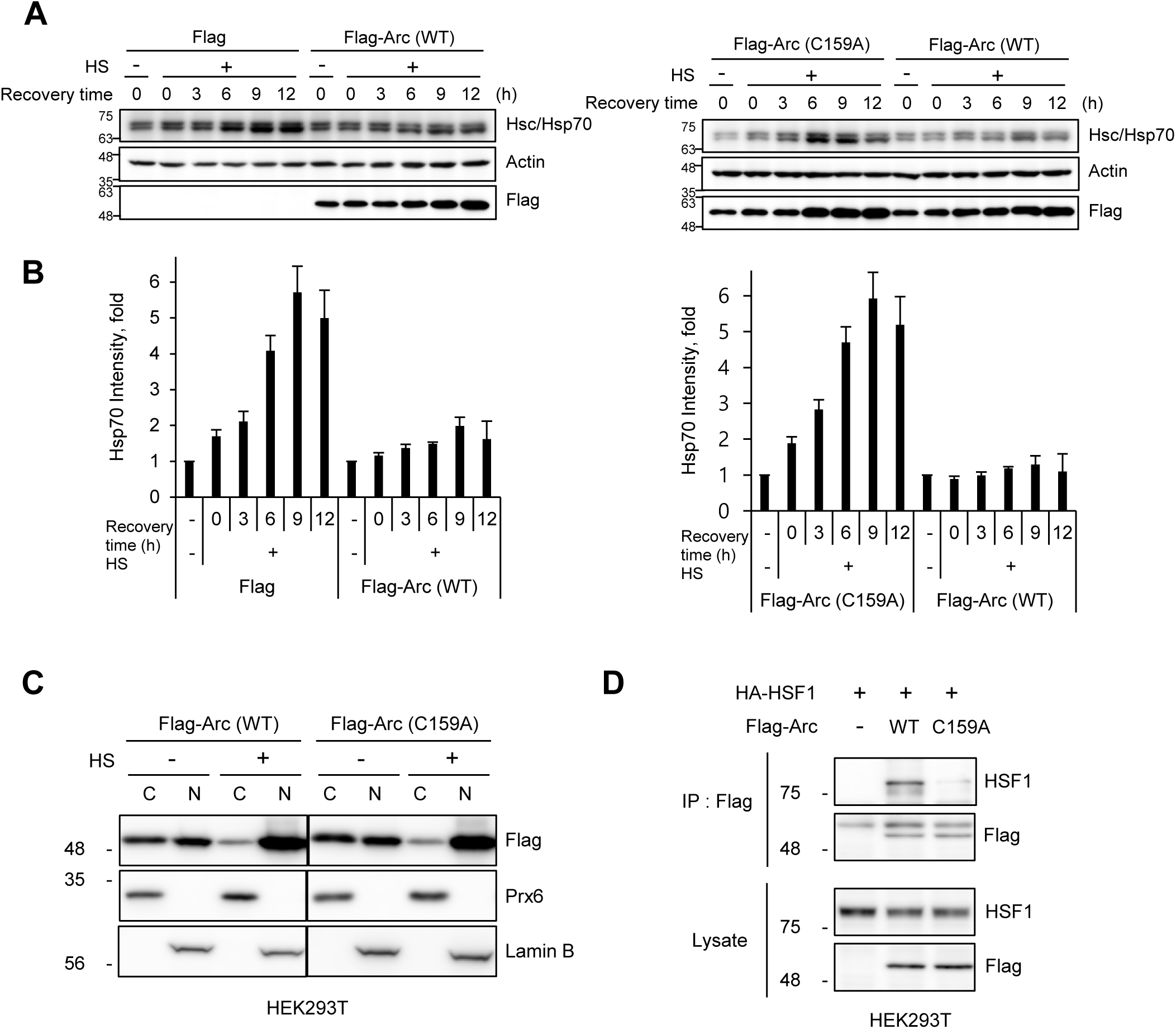
Arc/Arg3.1 negatively regulates the expression of HSP70 by binding to HSF1. (A, B) Flag-Arc(C159A) has no adverse effects on Hsp70 expression compared to Flag-Arc(WT). HEK293T cells overexpressing the Flag-Arc WT or C159A mutant were exposed to heat shock at 45 °C for 15 min and allowed to recover at 37 °C for the indicated times. Hsp70 and Flag-Arc/Arg3.1 were detected by Western blot analysis using their specific antibodies. Actin levels were detected by the anti-actin antibody and used as the loading control. Western blot results were selected as representative data from triplicated experiments. Quantified results of triplicated Western images are shown in (B). (C) Arc WT and C159A translocate into the nucleus after heat shock. HEK293T cells overexpressing Flag-Arc WT or the C159A mutant were exposed to heat shock at 45 °C for 15 min. Cytosolic and nuclear fractionation was performed. Each fraction was analyzed by Western blot analysis using anti-Flag, anti-Prx6, and anti-Lamin B antibodies. Prx6 is the cytosol marker; lamin B is the nucleus marker. (D) Only Arc WT, not C159A mutant, interacts with HSF1. HEK293T cells transfected with HA-HSF1 and Flag-Arc(WT) or Flag-Arc(C159A) were immunoprecipitated using an anti-Flag antibody. Western blot analysis used anti-HA and anti-Flag antibodies to analyze the protein complex.

### 2.9. Arc/Arg3.1 WT and C159A mutant each possess different interacting proteins

To investigate the further functional differences, we examined the interacting proteins of Arc/Arg3.1 WT and C159A mutant by employing immunoprecipitation and proteomic analysis. HEK293T cells were transfected with Flag empty vector, Flag-Arc WT, or Flag-Arc C159A, and the immune complexes precipitated by anti-Flag antibody were separated by SDS-PAGE and detected using silver staining (Fig. S6A). The most differentially appearing protein bands were identified by peptide sequencing with nanoUPLC-ESI-q-TOF tandem MS. The identified proteins of each band are listed in Table S5. Of these, COPS4, a component of the COP9 signalosome complex, was confirmed by Western blot (Fig. S6B). Arc/Arg3.1 interacting proteins were classified based on subcellular localization and biological process (Fig. S6C). The identified interacting proteins were matched with known functions of Arc/Arg3.1: Capsid interaction (cyclophilin), brain (Ataxin and Drebrin), membrane trafficking and endocytosis, and cytoskeleton. Arc/Arg3.1 is also associated with proteins related to synthesis and degradation of Arc/Arg3.1, including translation, mRNA decay (Upf1), ribosome, and UPS (COPS4). Chaperones (Hsp70s, Hsp27s, and Hsp90) are also identified, in which Hsp70 is verified to interact with Arc/Arg3.1 (Fig. 2D). Proteins related to the nucleus and nuclear translocation represented a nuclear function of Arc/Arg3.1, as suggested in a previous study (10). The results indicate that the aggregated and easily degradable Arc/Arg3.1 WT is required to interact with the membrane, nucleus, and mitochondrial and endocytosis proteins.

## 3. Discussion

The present study shows that Arc/Arg3.1, transiently expressed in response to heat shock stress and readily degraded, is not degraded in HSF1-/- MEF cells. We found that the expression of Hsp70 is required for the degradation of Arc/Arg3.1, which is caused by the E3 ligase CHIP interacting with Hsp70. Moreover, Arc/Arg3.1 oligomer and aggregate are formed through Cys34-Cys159 disulfide crosslinking, which was detected with MS/MS (38), NPSB-B labeling (47), and mutant studies. We found that Arc/Arg3.1 WT has a more closed structure and is readily aggregated by disulfide formation and degraded by the UPS. This study unveiled that the expression level of Arc/Arg3.1 is tightly regulated by negative feedback loops between Hsp70 and Arc/Arg3.1 because Hsp70 induced by HSF1 inhibits Arc/Arg3.1 activity by inducing oxidation-dependent degradation, while heat shock-induced Arc/Arg3.1 inhibits HSF1 activity (10).

This is the first report to explain the novel degradation pathway of Arc/Arg3.1 as a short-lived protein. Arc/Arg3.1 is induced by various stresses, including heat shock, such as Hsps, but is readily degraded by the UPS during recovery (t_1/2_, <30 min), unlike Hsps. Transiently expressed Arc/Arg3.1 inhibits *hsp* gene activation by blocking the binding of HSF1 to HSE. Thus, Arc/Arg3.1 is a novel regulator of activated HSF1 activity (10). This study shows that Hsp70 induced by HSF1 in response to heat shock causes Arc/Arg3.1 degradation by CHIP, an E3 ligase that interacts with the C-terminal of Hsp70 to form a negative feedback loop between Arc/Arg3.1 and Hsp70. Degradation of Arc/Arg3.1 by the UPS requires structural changes to oligomers and aggregates by forming Cys34-Cys159 disulfide. The Cys159 residue in Arc/Arg3.1, identified as a reactive cysteine residue, plays the critical regulatory function in forming the Cys34-Cys159 disulfide bond, which regulates the folding between the NTD and CTD in the Arc/Arg3.1 protein and results in the structural changes. The C159A mutant was not degraded by the UPS, which indicates that Arc/Arg3.1 is subject to redox regulation by disulfide formation with the reactive Cys159.

Since there is a flexible central linker between the two domains with opposite charges, a theoretical conformational change (open and closed state of monomer) of Arc/Arg3.1 was suggested by several combined experiments (54). However, in the recent structural analysis of WT Arc/Arg3.1, only the closed state, where the NTD interacts with CTD, followed by reduced linker region flexibility (39), was confirmed. In contrast, the weakening of the interaction between the NTD and CTD, shown in the Arc/Arg3.1 C159A mutant in this study by the HDX-MS experiment, indicates an open state by the Arc/Arg3.1 monomer. The disulfide bond formation by Arc/Arg3.1 may represent the mechanism that controls the transition from the open and closed structural states.

Recent studies have shown that the Arc/Arg3.1 dimer formation may be the first step towards capsid-like structure construction (39), whereby an Arc/Arg3.1 tetramer formation appears to depend on the N-terminal region of Arc/Arg3.1 (46). According to the oxidative capsid assembly mechanism of HIV-1, a budding virus enters the oxidizing environment of the bloodstream, where it forms disulfide bonds. These bonds can trigger CA oligomerization and subsequent maturation by regulating the interaction between capsids (55, 56). In addition, several studies of virus capsid assembly address the role of disulfide bonds in regulating capsid assembly and stability. Considering the life cycle of being released out of the cell to form a capsid structure, Arc/Arg3.1 is likely to follow the oxidative capsid assembly mechanism, similar to the HIV-1 virus. Since the Arc/Arg3.1 C159A or C34A mutants do not form oligomers or aggregates and are not degraded by the UPS in this study, dimer formation via disulfide crosslinking is possibly a key early stage for self-assembly of Arc/Arg3.1.

The structural and functional regulation of proteins via disulfide formation of redox-sensitive Cys residue appears as a key regulatory machinery of proteins. However, this mechanism has been neglected because proteins are generally handled in reducing DTT and β-ME conditions, meaning the disulfide information is lost. A recent study showed the methodology to identify the redox-sensitive Cys residues by employing a specific chemical probe NPSB-B for labeling redox-sensitive Cys residues (47) and found similar disulfide regulations in various proteins and Arc/Arg3.1.

By employing dynamic structural analyses using HDX-MS and functional studies, similar oxidative regulation in DJ-1(57), Nm23-H1 (NDPK-A) (52), peroxiredoxin 2 (Prx-2) (34), and human secretagogin (hSCGN) by Ca^2+^ binding (58) have also been elucidated. In DJ-1, Cys46-Cys53 intra-disulfide bonding with reactive Cys46, which was neglected in reducing conditions, was shown to be a crucial factor in the structural integrity and antioxidant activity of DJ-1 (57). Previous studies have concentrated on the function of Cys106 oxidation to sulfinate for its cytoprotective activity of DJ-1 (59). However, it turned out that the Cys106 oxidation state can be regulated by Cys46-Cys53 intra-disulfide bonding. Nm23-H1/NDPK-A, a tumor metastasis suppressor, is readily oxidized to form Cys4-Cys145 intra-disulfide crosslinking, which triggers significant structural changes from the hexamer and dimer and induces the loss of enzymatic activity by oxidizing Cys109 to sulfonic acid, similar to stepwise oxidation (50). Oxidized peroxiredoxin 2 (Prx2) is readily degraded in the proteasome, not reduced. It turns out that oxidation-induced structural change makes the protein ubiquitinated in redox-sensitive proteins (34). Secretagogin, a hexa EF-hand calcium-binding protein that plays an essential role in insulin secretion in pancreatic beta-cells, induces structural change through calcium binding, which induces functional dimer formation via an inter-disulfide bond(58). To prove this concept, we investigated the structural changes by the Arc/Arg3.1 WT and C159A mutant using HDX-MS analysis and structure modeling. Here, we found that the aggregation properties of the Arc/Arg3.1 WT were based on a close conformation between the NTD and CTD. Conversely, the C159 mutant, which does not form an intra-disulfide bond, has an open structure by attenuating the interaction between the NTD and CTD. In summary, this study identifies the degradation mechanism for Arc/Arg3.1, with the shortest half-life (< 30 min), via disulfide formation to form oligomers and aggregates. Moreover, the expression level of Arc/Arg3.1 is tightly regulated by Hsp70 expression via the activation of CHIP, an E3 ligase that interacts with Hsp70. Various stresses induce Arc/Arg3.1 and activate HSF1; overexpressed Arc/Arg3.1 impeded activated HSF1, and Hsp70 expressed by activated HSF1 expedites the degradation of Arc/Arg3.1 through a negative feedback loop. Whether this molecular mechanism of Arc/Arg3.1 regulation is necessary for its diverse functions can only be answered by further studies.

## 4. Materials and Methods

Detailed methods are found in supplemental methods section

### 4.1. Cell culture and heat shock treatment

WT and HSF1-/- MEF cell lines (gifts from Dr. Ivor J. Benjamin, University of Utah, USA) were grown in high glucose Dulbecco’s Modified Eagle Medium (DMEM) supplemented with 10% fetal bovine serum (FBS), 100 units/mL penicillin G and 100 /mL streptomycin at 37 °C in a 5% CO_2_-containing humidified incubator. HEK293T and HeLa cells were purchased from ATCC and maintained following the manufacturer’s protocol. Radiation-induced mouse fibrosarcoma cell line, RIF-1, was grown in RPMI1640, supplemented with 10% fetal bovine serum (FBS), 100 μg/mL streptomycin, 100 units/mL penicillin G, 3.75 μg/mL sodium bicarbonate, and 0.11 μg/mL sodium pyruvate at 37 °C in a 5% CO_2_-containing humidified incubator. Heat shock was administered to cells grown in tissue culture dishes by placing them in a water bath at 45 °C.

### 4.2. Protein expression and purification

A pET-28a(+) plasmid carrying the His-hArc/Arg3.1 gene was expressed in BL21 (DE3) *E.coli* cells. Cells were grown in an LB medium containing 50 mg/L kanamycin at 37 °C, with shaking. At an optical density (OD) of 0.7, 0.2 mM isopropyl-β-D-thiogalactopyranoside (IPTG) was added. After induction for 4 h, cell pellets were harvested by centrifugation, resuspended by vortexing with PBS buffer containing phenylmethylsulfonyl fluoride (PMSF, 0.1 mM), 2 μg/mL aprotinin, 20 μM leupeptin, 10 μg/mL pepstatin A, 1 mM DTT, 1.5% (w/v) N-lauryl sarcosine, and 10 mM imidazole and . 100 μg/mL of lysozyme was added to the cell suspension. After being sonicated, 2% (w/v) Triton X-100 was added to the cell lysates and remained on ice for 30 min with vortexing at 5 min intervals. After centrifuging, the supernatants were applied to Ni-NTA agarose beads (Qiagen, USA) at 4 °C for 4 h. The beads were washed with 10 mL phosphate buffer in saline with 0.1% Triton X-100 (PBST) containing 0.5 M NaCl and 10 mM imidazole, 10 mL PBST containing 30 mM imidazole, and 10 mL PBS containing 50 mM imidazole. Purified Arc/Arg3.1 was eluted with PBS containing 200 mM imidazole.

### 4.3. Disulfide analysis of hArc/Arg3.1 using the nanoUPLC-ESI-q-TOF tandem MS and DBond algorithm

The gel bands separated on the non-reducing condition without β-ME were destained and digested with trypsin. The extracted peptides were sequenced by nanoAcquity^TM^ UPLC^TM^/ESI/MS (SYNAPT^TM^ HDMS^TM^, Waters Co. UK) and analyzed by the disulfide searching algorithm DBond (http://prix.hanyang.ac.kr) (38).

### 4.4. Protein structure modeling

The Robetta online server (http://new.robetta.org) was used to generate a full-length human Arc/Arg3.1 model. The hArc/Arg3.1 sequence was submitted to the server and parsed into putative domains. The structural model was generated using comparative modeling methods.

### 4.5. Identification of reactive cysteine residues employing NPSB-B labeling

As previously described, reactive cysteine residues can be detected by specific chemical probe NPSB-B labeling (47). Recombinant hArc/Arg3.1 protein (10 μM) purified with DTT was labeled with 1 mM NPSB-B at RT for 2 h. Recombinant hArc/Arg3.1 protein (5 μM) was treated with indicated concentrations of H_2_O_2_ at 37 °C for 1 h before being labeled with 1 mM NPSB-B at RT for 2 h. Labeled proteins in the SDS gel sample buffer containing 20 mM NEM were detected using streptavidin-HRP. The amount of loaded protein was detected by coomassie blue staining.

### 4.6. Hydrogen/deuterium exchange mass spectrometry (HDX-MS)

Arc/Arg3.1 WT and C159A proteins (1 μg/μL) were diluted 10-fold with labeling buffer (20 mM PBS in D_2_O, pH 7.4) and maintained at 25 °C for several time scales. The labeling reaction was quenched by 25 mM tris (2-carboxyethyl) phosphine (TCEP) buffer, pH 2.3. For peptic digestion, porcine pepsin (1 μg) was added to each quenched protein sample and incubated at 0 °C for 3 min before injection. The autosampler chamber was set at 4 °C. The trap, analytical column, and all tubing were immersed in an ice bath to minimize deuterium back-exchange. The gradient chromatography was performed at a flow rate of 0.52 μL/min and was sprayed online to nanoAcquity^TM^/ESI/MS equipped at Ewha Drug Development Research Core Center (SYNAP^TM^ HDMS^TM^, Waters, Co.). All mass spectral measurements were taken at capillary voltage 2.5 kV, cone voltage 35 V, extraction cone voltage 4.0 V, and source temperature 80 °C. TOF mode scan was performed at the m/z 300-1500 range with a scan time of 1 s.

## Acknowledgements

We are grateful to Dr. Ivor J. Benjamin (University of Utah, USA) for donating HSF-/- MEF cell line. The authors appreciate the technical support of the MS analysis performed by Mr. Kang W.

## Author contributions

D.S, I.-K.S., Y.J.K., H.-J.K., K.-J.L., and E.J.S. designed research; D.S., and Y.J.K. performed biochemical research; I.-K.S. and Y.S.P. performed mass spectrometry; Y.L. performed DLS; D.S, I.-K.S., Y.J.K., H.-J.K., K.-J.L., and E.J.S. analyzed data; D.S, I.-K.S., and Y.J.K. wrote the paper. H.-J.K., K.-J.L., and E.J.S. supervised the research.

## Data availability

All data needed to evaluate the conclusions in the paper are present in the paper and/or the Supplementary Materials. The structures presented in this paper have all been deposited in the Protein Data Bank (PDB) with the following codes: rArc/Arg3.1 N-lobe (PDB: 4X3H) and rArc/Arg3.1 C-lobe (PDB: 4X3X). Raw Arc/Arg3.1 DBond and HDX analysis data (Fig. 4B, 7, Table 4) are available via ProteomeXchange with the identifier PXD026534 (60). Reviewer account details are as follows; password: K26AjDHl.

---

**Supplementary Figure S1.**
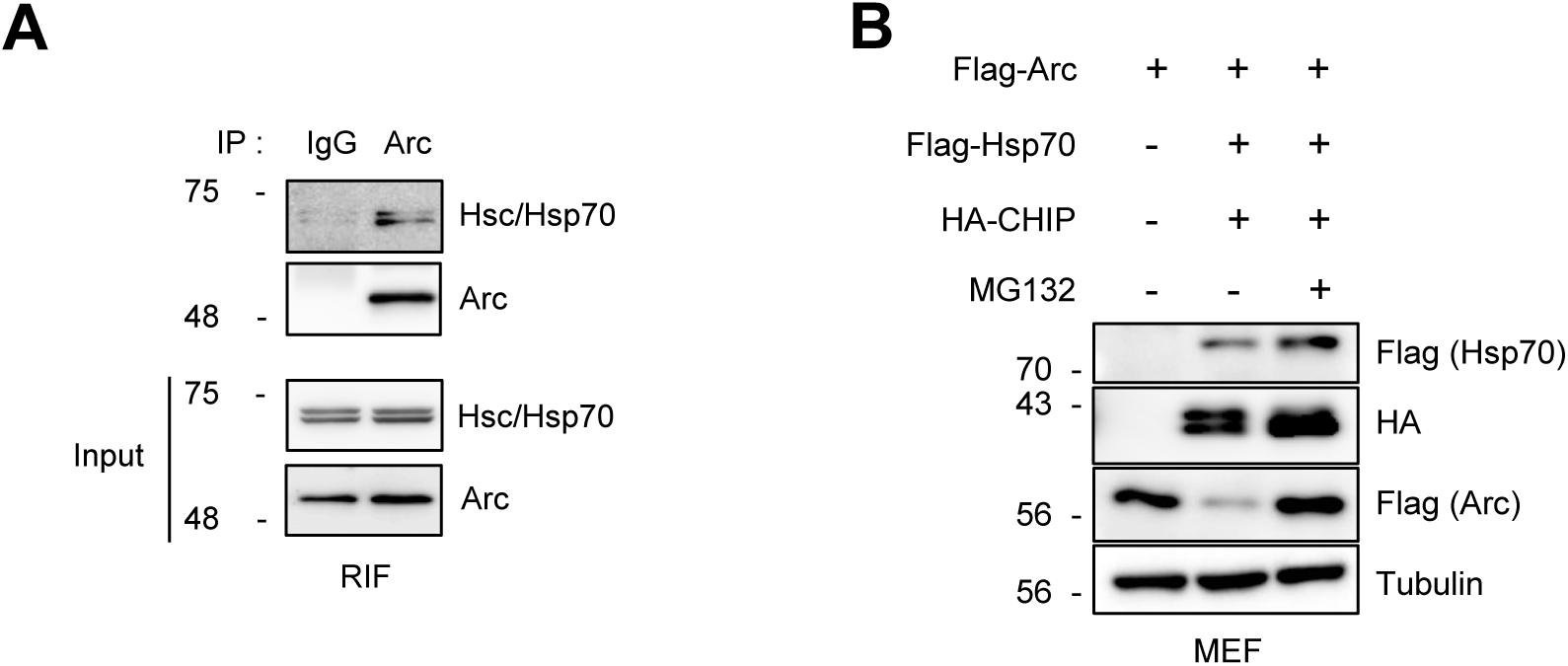
(A) Interaction of endogenous Arc/Arg3.1 and Hsp70. RIF cells were exposed to heat shock (45°C for 25 min) and recovered for 6 h at 37°C. Cell lysates were immunoprecipitated using anti-Arc antibody. Immune-complex was analyzed by Western blot with anti-Arc and anti-Hsc/Hsp70 antibodies. (B) The restoration of Arc/Arg3.1 proteins upon MG132 treatment. After being transfected with Flag-Arc for 24 h, either in the presence or absence of both Flag-HSP70 and HA-CHIP, MEF cells were exposed to 20 μM MG132 for a duration of 6 h.

**Supplementary Figure S2.**
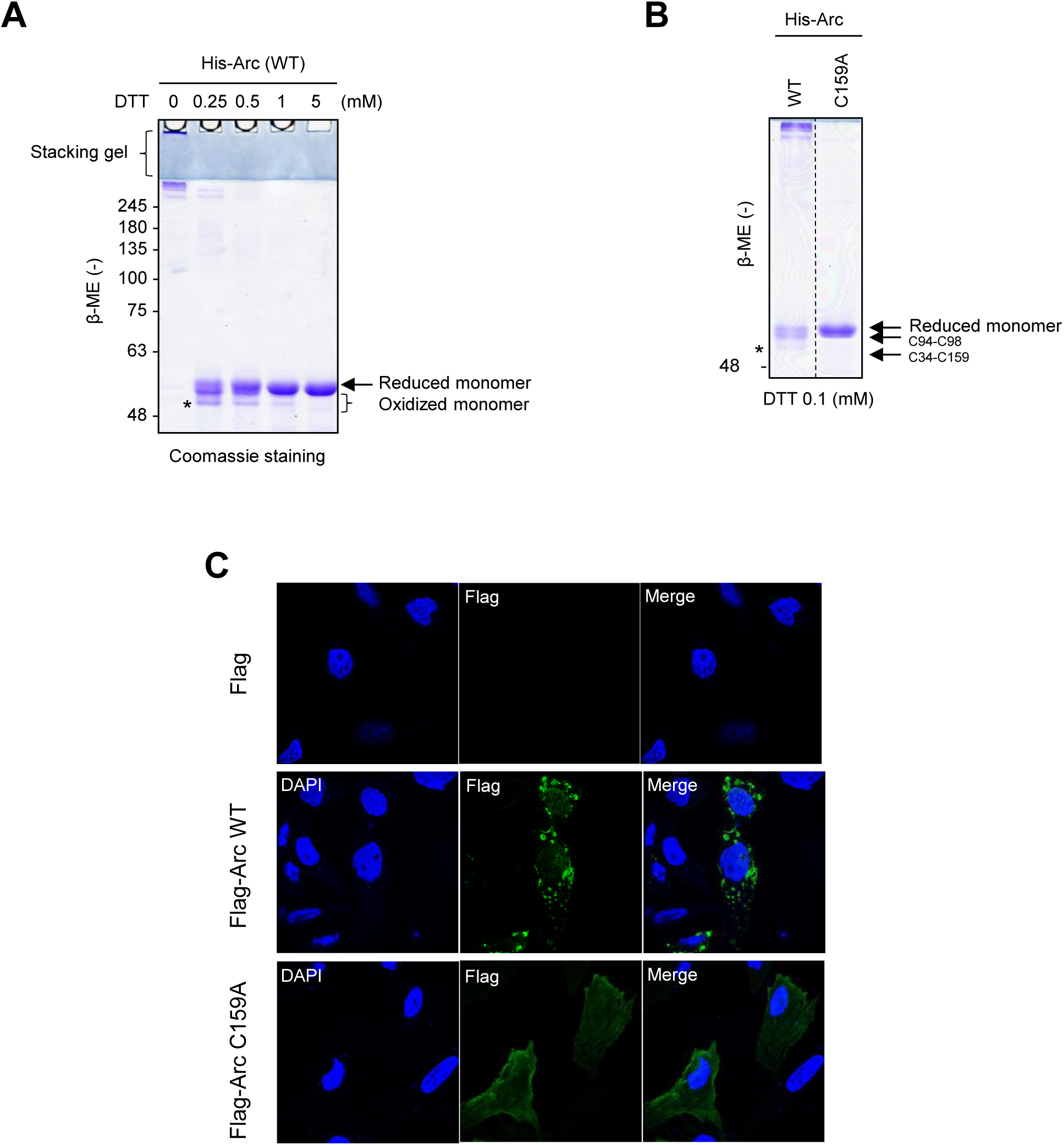
(A) Purified hArc/Arg3.1 protein was treated with indicated concentration of DTT at 37°C for 30 min. Proteins were separated by SDS-PAGE under non-reducing (-β-ME) conditions and Coomassie blue stained. (B) Purified hArc/Arg3.1 WT and C159A proteins were treated with 0.1 mM DTT at 37°C for 30 min. Proteins were separated by SDS-PAGE under non-reducing (-β-ME) conditions and Coomassie blue stained. (C) For exogenous Arc/Arg3.1, cells were transfected with Flag, Flag-Arc WT or Flag-C159A. Cells were fixed and stained for Flag-Arc (green). DAPI (blue) was used for nucleus staining. Cells were photographed at x 40 magnification with a fluorescence confocal microscope.

**Supplementary Figure S3.**
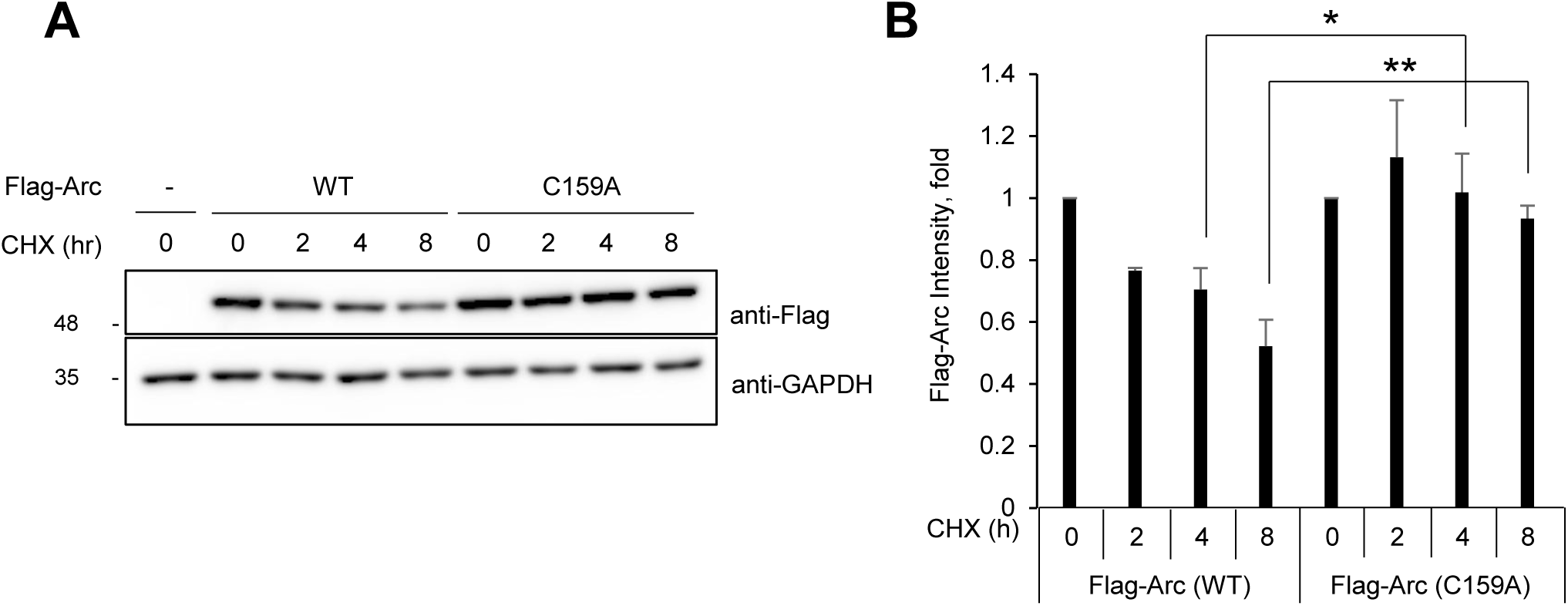
(A) HEK293T cells overexpressing Flag-Arc WT or C159A mutant were treated with 10 μg/mL cycloheximide (CHX) for the indicated times after the treatment of 0.25 mM H_2_O_2_ at 37°C for 1 h. The Arc/Arg3.1 protein degradation was analyzed by Western blotting using an anti-Flag antibody. GAPDH was used as a loading control. Representative Western results were selected from triplicated experiments. (B) Quantified results of quadruplicated Western images are shown in (A).

**Supplementary Figure S4.**
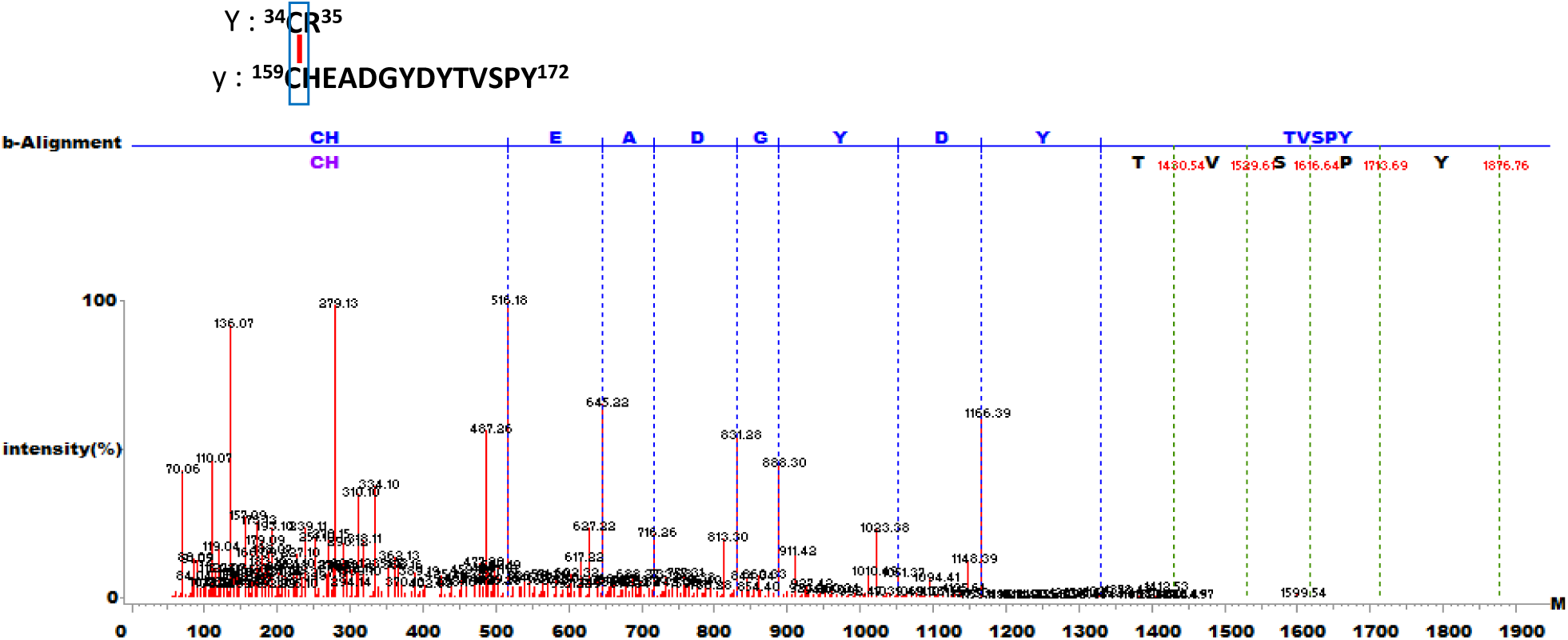
Tandem mass spectrum of Cys34-Cys159 disulfide bond peptide in the WT in Fig. 4B. Coomassie-stained gels were cut out of the gel with trypsin and chymotrypsin and analyzed by MS and Dbond analysis was performed.

**Supplementary Figure S5.**
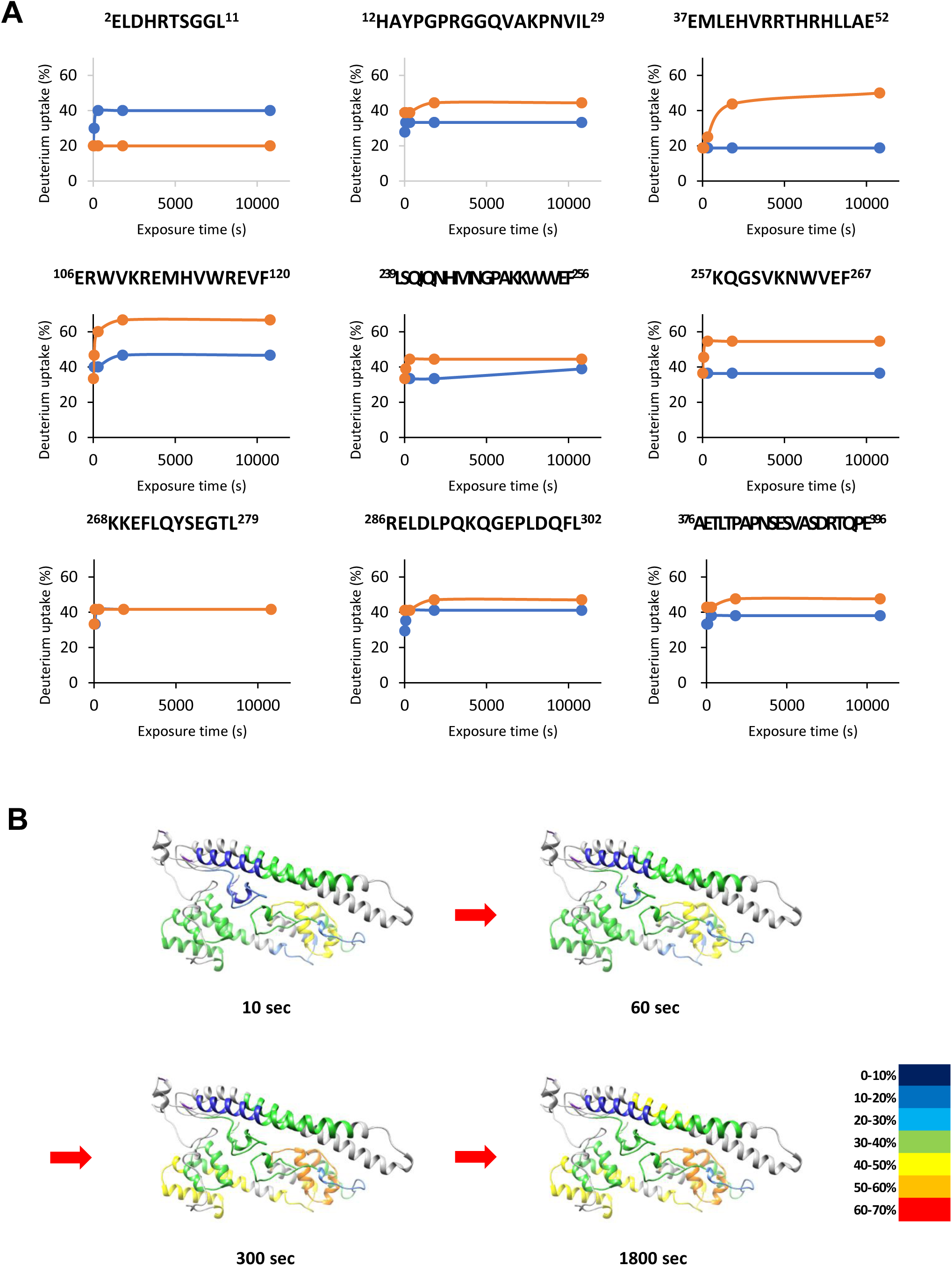
(A) Time course of HDX incorporation for representative peptides that showed differences in HDX(WT: blue, C159A: red). (B) Changes in HDX-MS of Arc/Arg3.1 WT as compared with the 0 s incubation at various incubation times.

**Supplementary Figure S6.**
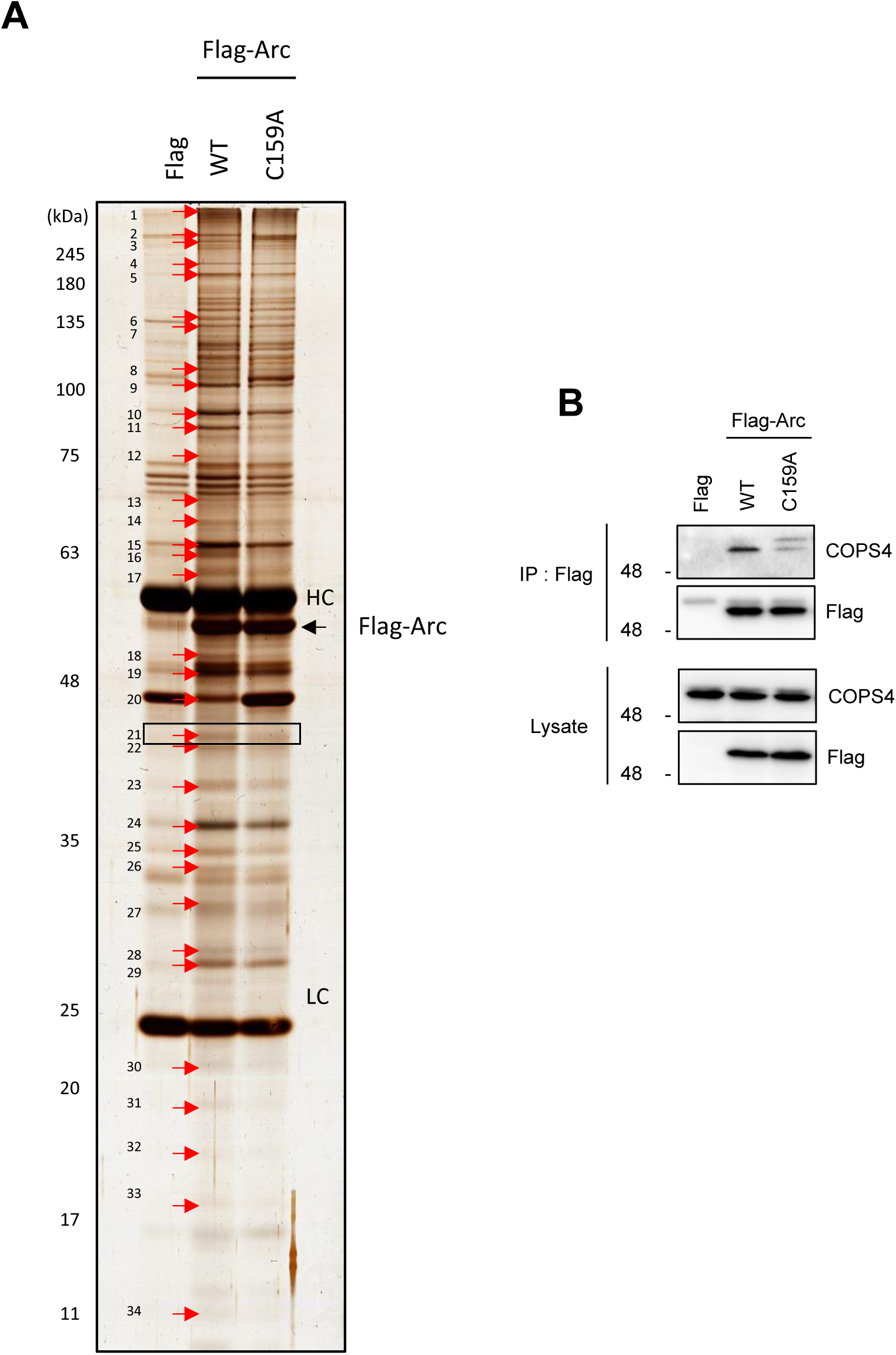
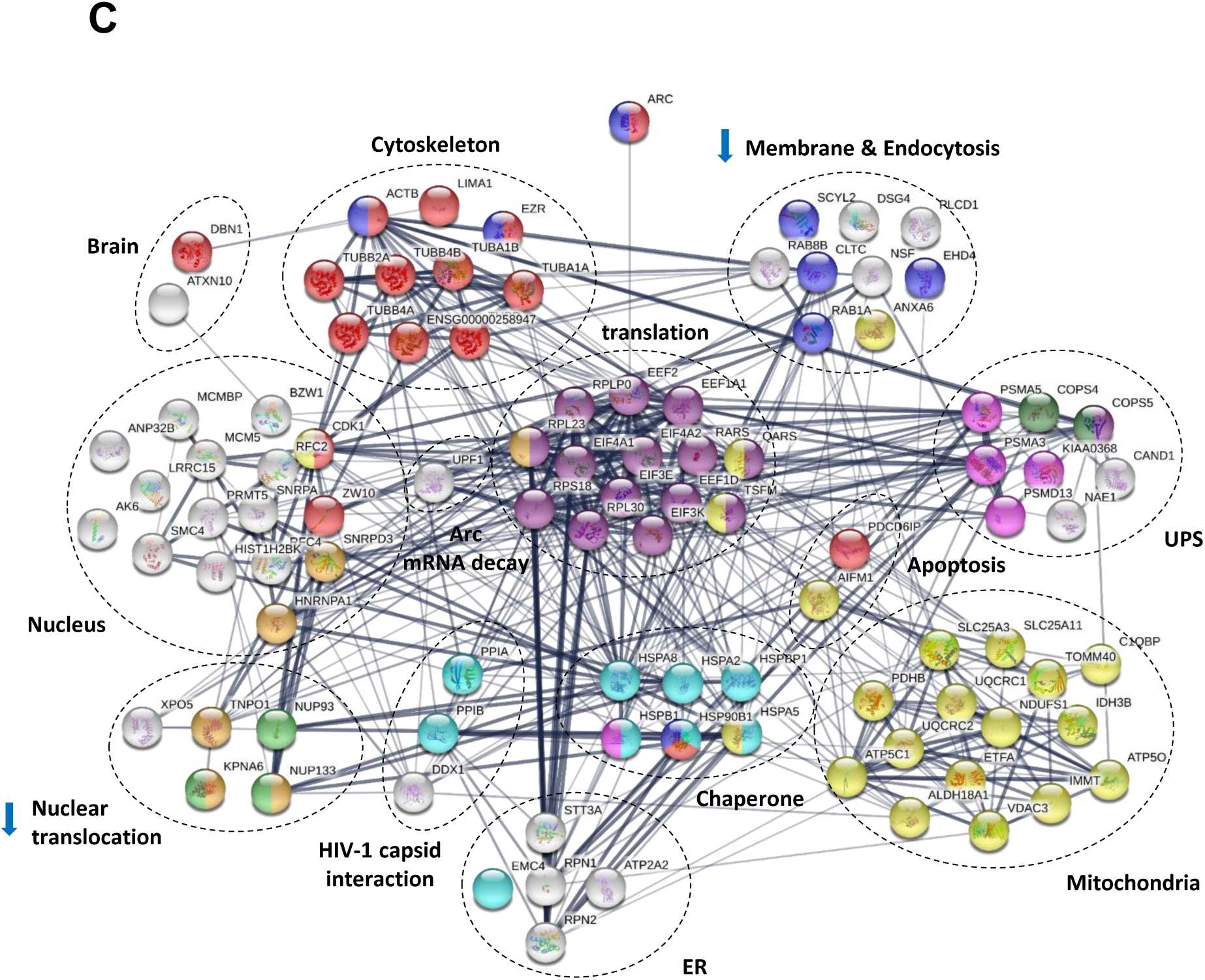
Identification of Arc/Arg3.1 interacting protein. (A) HEK293T cells were transfected with Flag empty vector, Flag-Arc WT or C159A. Immunoprecipitation was performed using anti-Flag antibody. The location of differential band subjected to further MS identification was indicated with number. (B) Interaction between Arc and COPS4 was confirmed by western blot analysis. (C) STRING protein network analysis represents Arc/Arg3.1 interacting proteins classified in Supplementary Table 5 (except miscellaneous). Edges represent protein-protein associations with confidence coefficient higher than 0.4. The thickness of the edge indicates the degree of confidence coefficient.

**Supplementary Table S1.**
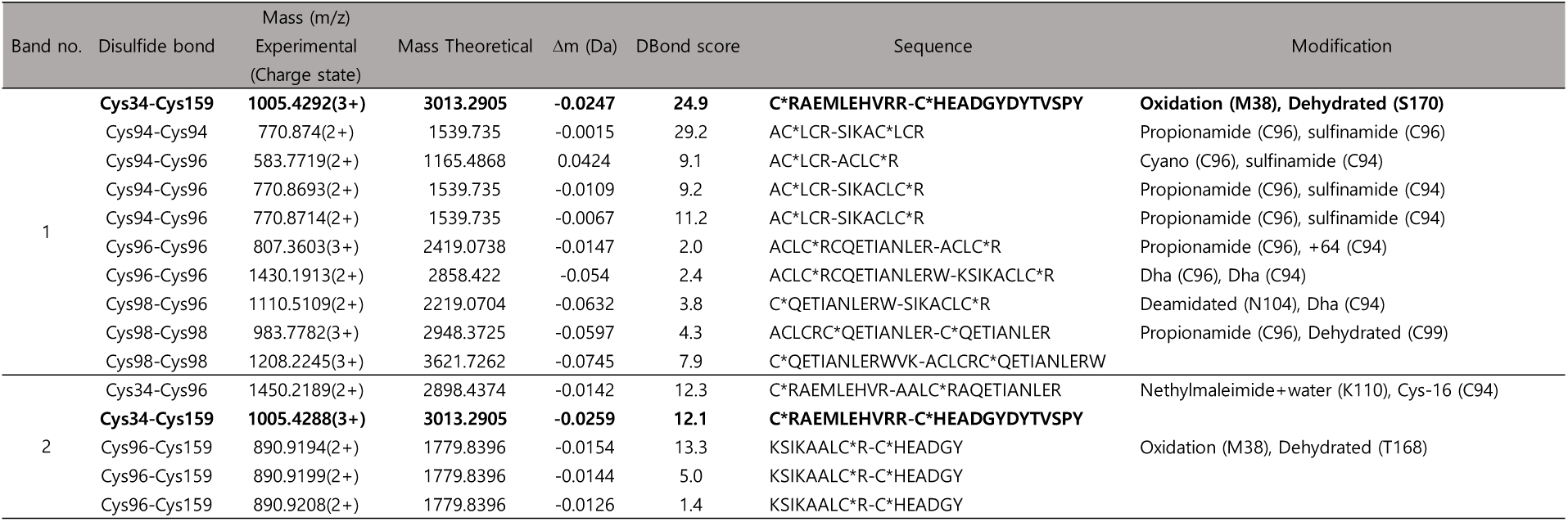
List of identified disulfide bonds in Arc WT and C94,98A mutant proteins (Fig. 4B) under the non-reducing condition. Protein were treated with NEM to avoid additional oxidation and separated non-reducing SDS-PAGE, and then analyzed with MS/MS combining with DBond algorithm.

**Supplementary Table S2.**
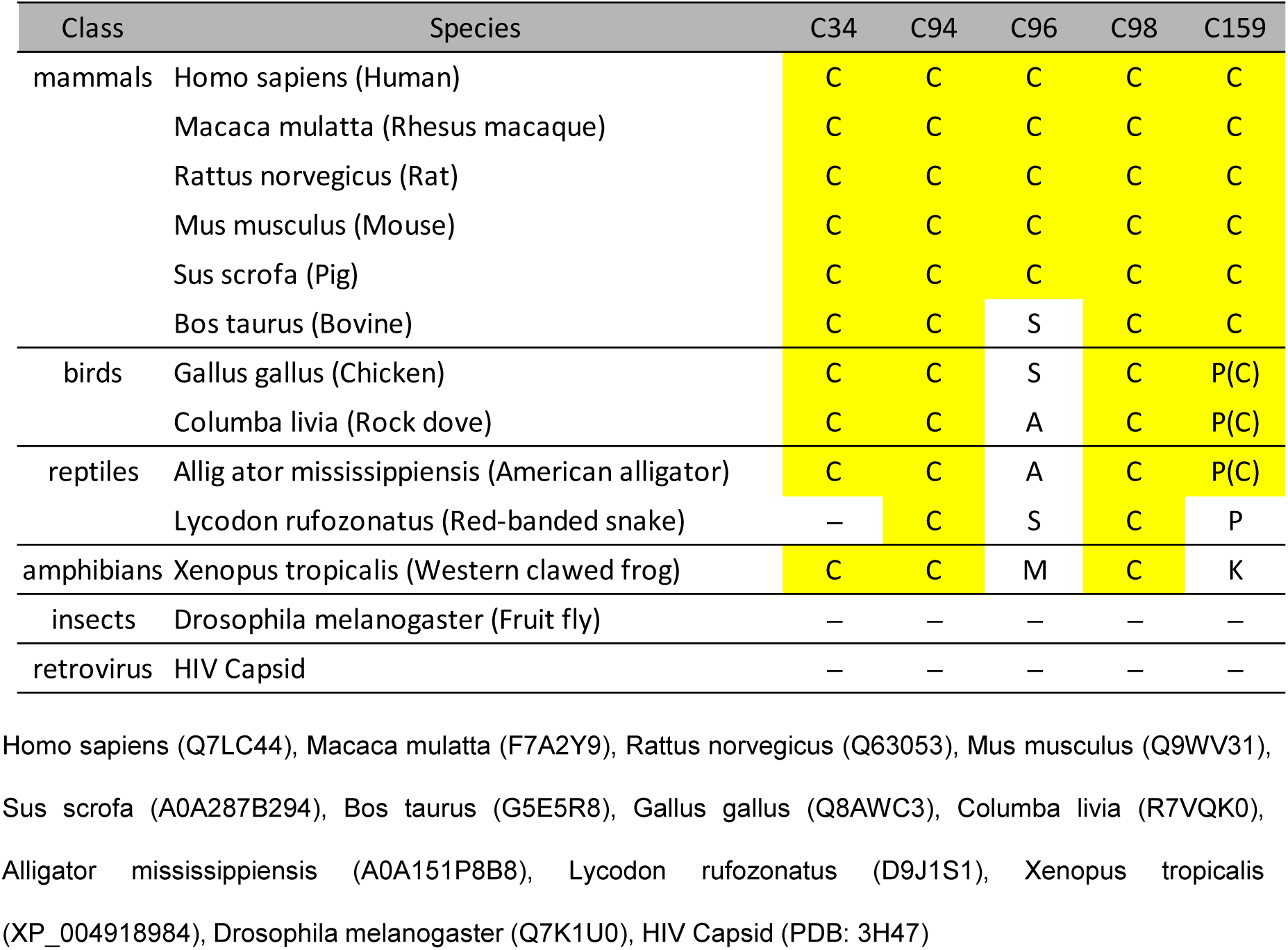
Evolution of Arc cysteines was studied by sequence alignment. The five mammalian cysteines in Arc show different evolutionary patterns

**Supplementary Table S3.** HDX-MS summary data

| Data Set | WILDTYPE | MUTANT |
| --- | --- | --- |
| HDX reaction details | 140 mM NaCl, 2.7 mM KCl, 10 mM<br>$\text{Na}_2\text{HPO}_4$, $\text{pD}_{\text{read}} = 7.4$, 25°C | 140 mM NaCl, 2.7 mM KCl, 10 mM<br>$\text{Na}_2\text{HPO}_4$, $\text{pD}_{\text{read}} = 7.4$, 25°C |
| HDX time course (min) | 0.1, 1, 5, 30, 180 | 0.1, 1, 5, 30, 180 |
| HDX control samples | Deuterium-non-labeled control (WT) | Deuterium-non-labeled control (mutant) |
| # of Peptides | 19 | 23 |
| Sequence coverage | 71% | 84% |
| Average peptide length | 15.68 | 16.09 |
| Replicates (biological or technical) | 2 (technical) | 2 (technical) |

**Supplementary Table S4.**
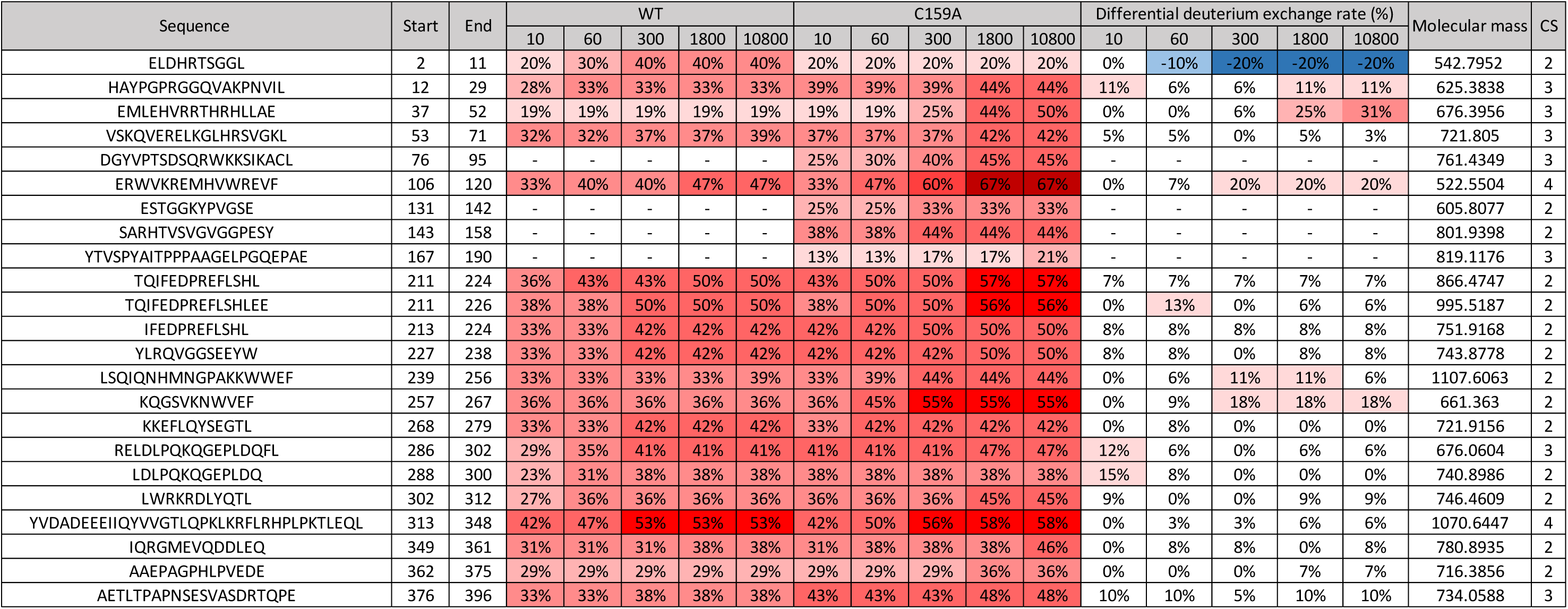
Differential deuterium exchange rates of identified Arc peptides in HDX-MS experiments.

**Supplementary Table S5.**
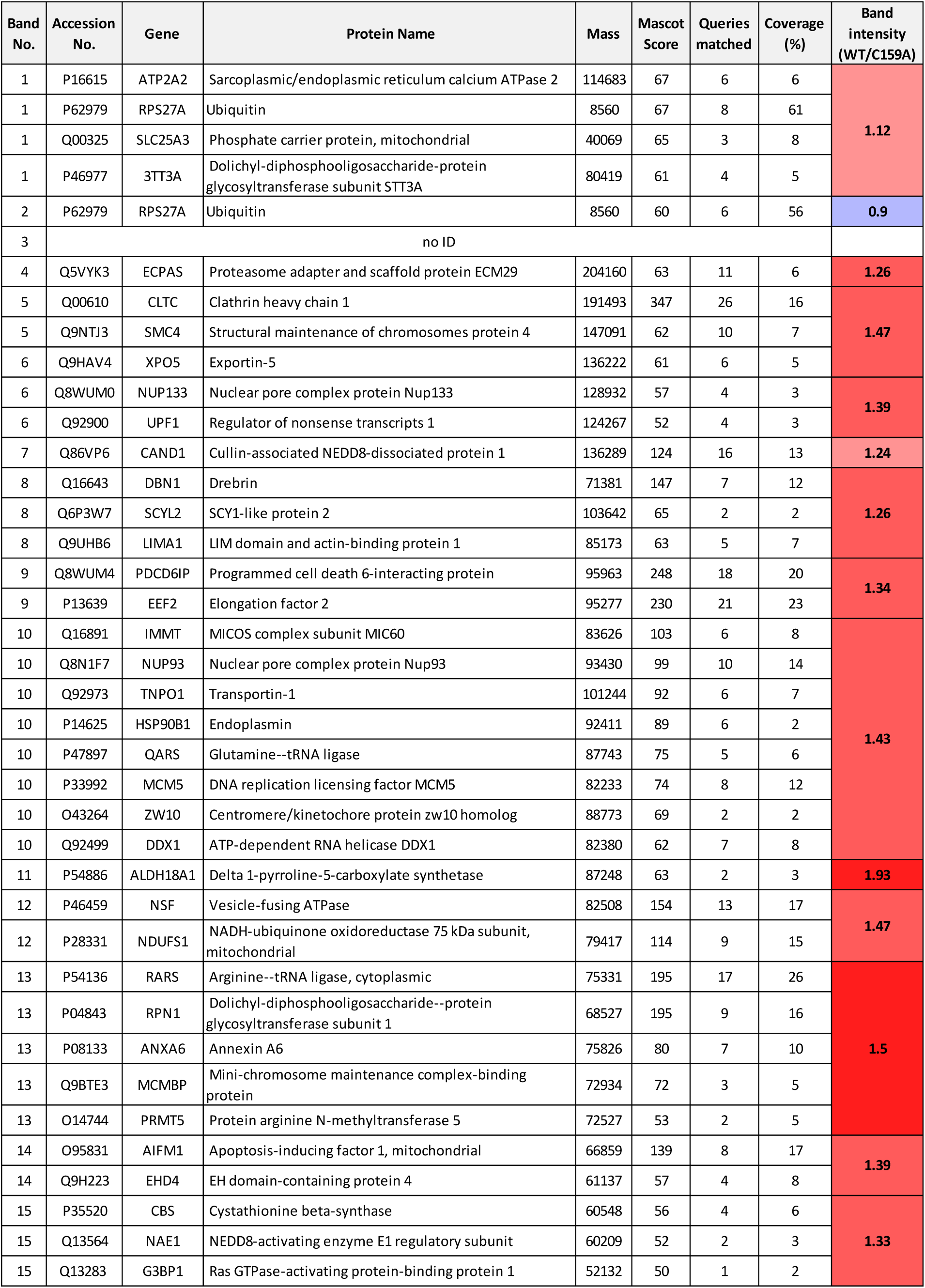
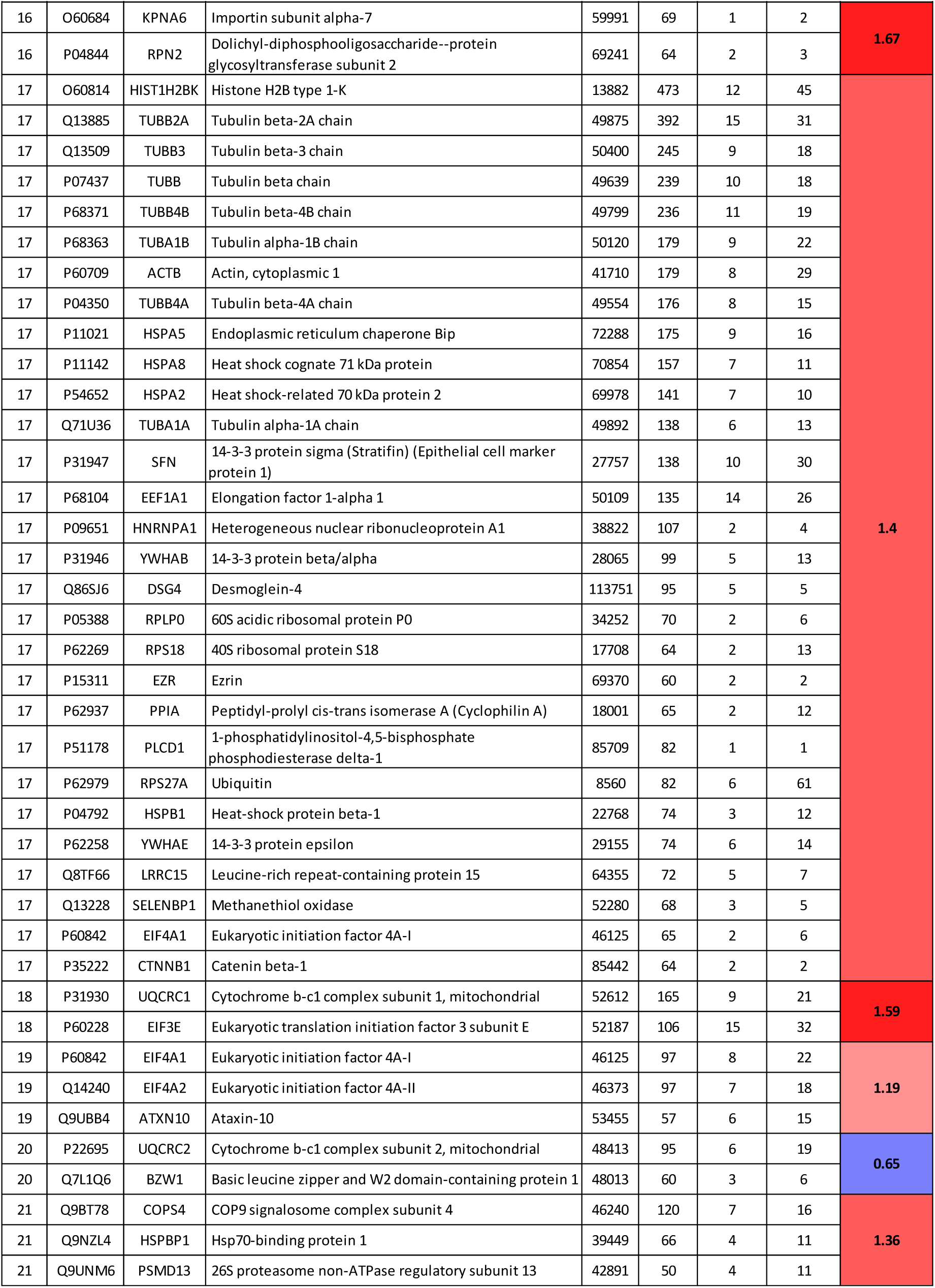
List of Arc binding proteins identified in each band. Each of thirty-four protein bands from Supplementary Fig. 6A was identified using nano-UPLC-ESI-q-TOF tandem mass spectrometry, classified manually based on subcellular localization and biological process.

## Supplemental Methods

### Antibodies and reagents

Mouse monoclonal Arc antibody (E-7) (sc-55475), mouse monoclonal Arc antibody (C-7) (sc-17839), mouse monoclonal anti-β-Actin (C4) (sc-47778), mouse monoclonal HSP27 antibody (sc-13132), and mouse monoclonal α-tubulin antibody (sc-8035) were purchased from Santa Cruz: Rabbit polyclonal GAPDH antibody (LF-PA0018) and mouse monoclonal HA antibody (LF7H5) (LF-MA0048) from Ab Frontier: Mouse monoclonal Hsc/Hsp70 antibody (ADI-SPA-820) and mouse monoclonal HSF1 antibody (ADI-SPA-901) from Enzo: Mouse monoclonal FLAG antibody (F3165) from Sigma Aldrich: Mouse monoclonal GFP antibody (A-11120) from Invitrogen: Streptavidin-HRP (3999), rabbit monoclonal BiP antibody (C50B12) (3177), and rabbit polyclonal Hsp40 antibody (4868) from Cell signaling: Mouse monoclonal Hsp90 antibody (ab13492), rabbit polyclonal Hsp60 antibody (ab13492), and rabbit polyclonal Hsp60 antibody (ab46798) from Abcam: Goat mouse IgG-HRP conjugate antibody (170–5047) and goat rabbit IgG-HRP conjugate antibody (170–6515) from Bio-Rad. Hydrogen peroxide was purchased from Samchun Pure Chemical Co. N-ethylmaleimide (E1271), cycloheximide (C7698) was acquired from Sigma Aldrich, and MG132 (M-1157) was procured from AG Scientific.

### DNA constructs (plasmids, cloning, and mutagenesis)

Human Arc/Arg3.1 was subcloned into the pFlag-CMV-2 vector and pET-28a(+) vector. For cloning the Flag-hArc/Arg3.1 DNA, human Arc/Arg3.1 cDNA were prepared by PCR using the sense primer 5’-GGA ATT CGA GCT GGA CCA CCG G-3’ and antisense primer 5’-CGG GAT CCC TAC TCG GGC TGG GT-3’. For cloning of the His-hArc/Arg3.1 DNA, PCR was performed using the same sense primer described above and antisense primer 5’-GCG GCC GCC TAC TCG GGC TGG GTC CGG TCA-3’. The PCR products and vectors were digested using EcoRI/BamHI and EcoRI/NotI and were ligated using the pGEM T-Easy vector system (Promega).

GFP-Hsp70 plasmid was a gift from Dr. Yun Sil Lee (Ewha Womans University, Korea). Flag-Hsp70 plasmid was a gift from Dr. Eunhee Kim (Chungnam University, Korea). HA-CHIP WT, H260Q, and K30A mutant plasmids were kindly provided by Prof. Jaewhan Song (Yonsei University, Republic of Korea).

Mutant hArc/Arg3.1 C34A, C94/98A, and C159A were generated using the Quik-Change^®^II Site-Directed Mutagenesis kit (Agilent Technologies), according to the manufacturer’s protocol using either pFlag-CMV-2-hArc/Arg3.1 or pET-28a(+)-hArc/Arg3.1 as a template. The mutagenesis oligonucleotides used were as follows: For mutation C34A, sense primer 5’-GCA GAT CGG GAA GGC CCG GGC CGA GAT G-3’, antisense primer 5’-CAT CTC GGC CCG GGC CTT CCC GAT CTG C-3’; for mutation C94/98A, sense primer 5’-TCC ATC AAG GCC GCC CTG TGC CGC GCC CAG GAG ACC AT-3’, antisense primer 5’-ATG GTC TCC TGG GCG CGG CAC AGG GCG GCC TTG ATG GA-3’; for mutation C159A, sense primer 5’-GTC CCG AGA GCT ACG CCC ACG AGG CAG ACG-3’, antisense primer 5’-CGT CTG CCT CGT GGG CGT AGC TCT CGG GAC-3’. All plasmid constructs were confirmed by DNA sequencing.

### Transient transfection

Cells were plated in a 35 mm culture plate for 24 h before transfection and treated with the indicated plasmid and TransIT-LT1 (Mirus, USA) reagent diluted in Opti-MEM (Gibco Life Technologies, USA), according to the manufacturer’s protocol. Cells were incubated with a DNA-transfection reagent complex for another 24-48 h before being used for further analysis.

### Dynamic light scattering (DLS)

Malvern Zetasizer Nano ZS90 was used for DLS experiments of purified Arc/Arg3.1 WT protein. The proteins were diluted to 0.5 mg/ml and the buffer was replaced with the indicated buffers using Amicon Ultra 0.5 mL Centrifugal filters (Merck Millipore) following the manufacturer’s instructions. Additional 1 mM dithiothreitol (DTT), 10 mM β-mercaptoethanol (β-ME), 0.5 mM sodium dodecyl sulfate (SDS) or H_2_O_2_ were added according to conditions. The proteins were equilibrated at the designated temperatures for 120 sec. The sample was measured at least twice, and the average of each replicate was calculated from 12 runs.

### Degradation of Arc/Arg3.1 by E3 ligases CHIP

To detect the degradation of Arc/Arg3.1 by the CHIP E3-ligase, HEK293T cells overexpressing Flag-Arc WT were co-transfected with His-ubiquitin and HA-CHIP WT, or H260Q, K30A mutant. Cells were harvested with an SDS-PAGE sample buffer and boiled at 95 °C for 10 min to elute the immune complexes. The samples were analyzed by Western immunoblotting.

### His-ubiquitin pull-down assay

For ubiquitination of Arc/Arg3.1, HEK293T cells overexpressing Flag-Arc WT or C159A were co-transfected with His-ubiquitin (His-Ub) and HA-CHIP. The appropriate plasmids were co-transfected with His-ubiquitin in HEK293T cells. The cells were resuspended in urea lysis buffer (pH 8.0, 8 M Urea, 0.3 M NaCl, 0.5 M Na2HPO4, 0.05 M Tris, 0.001 M phenylmethylsulfonyl fluoride (PMSF), 0.01 M imidazole) following 48 h transfection period, and sonicated for 2 min. Following the addition of cell lysates, 100 µL of equilibrated Ni-NTA agarose (Qiagen) was incubated for 4 h.

### SDS-PAGE and Western blot analysis

Whole-cell lysates were separated by SDS-PAGE and transferred to a polyvinyl difluoride (PVDF) membrane. The membranes were blocked with 5% BSA in PBS for 1-2 h and sequentially incubated with each antibody diluted in PBST at 4°C overnight, according to the manufacturer’s instructions (PBS buffer containing 3% BSA and 0.1% Tween 20). After washing three times with PBST for 10 min, the membrane was incubated with a secondary antibody, diluted in PBST to 1:3000-1:5000, incubated at RT for 45 min, and then rewashed three times. Amersham ECL Prime Western blotting detection reagent (GE Healthcare, UK) and an Amersham Imager 600 (GE Healthcare, UK) were used to detect the chemiluminescent signals, and Multi Gauge V3.0 (Fujifilm, Japan) was used for image analysis.

### Immunoprecipitation

After HEK293T cells were overexpressed with the plasmids, cells were harvested cells after 24 h and lysed in IP buffer (50 mM Tris-Cl, 150 mM NaCl, 1 mM EDTA, 0.5% Nonidet P-40, 60 mM octyl-β-D-glucopyranoside, pH 7.4) supplemented with protease inhibitor cocktail (Sigma-Aldrich, USA), phosphatase inhibitors (5 mM Na_3_VO_4_ and 5 mM NaF), and histone deacetylase inhibitors (10 mM sodium butyrate and 10 μM Trichostatin A). Cells were passed a 31G syringe 10 times, incubated on ice for 30 min, and then centrifuged at 12,000 rpm for 1 h. The supernatant was incubated with an anti-Flag antibody at 4 °C for 2 h. The lysate-antibody complexes were incubated with protein-G sepharose 4 Fast Flow beads at 4 °C for another 1 h. The precipitated beads were washed five times with washing buffer (50 mM Tris-Cl, 150 mM NaCl, 1 mM EDTA, and 0.5% NP-40, pH 7.4) and three times more with washing buffer without detergent. Immune complexes were separated by SDS-PAGE and detected using Western blot analysis and silver staining.

### Differential detergent extraction

HEK293T cells were lysed in RIPA buffer (50 mM Tris-HCl, pH 7.4, 150 mM NaCl, 1 mM EDTA, 1% (v/v) NP40, 0.5% (w/v) sodium deoxycholate, and 0.1% (w/v) SDS) supplemented with protease inhibitor cocktail (Sigma). Cell lysates were passed through a 31G syringe 10 times and centrifuged at 12,000 rpm for 20 min. The supernatant (soluble fraction) was separated, and the pellet (insoluble fraction) was washed twice with lysis buffer.

### Confocal microscopy

Cells were grown on the secureslip™ (Gracebiolab, OR, USA) cell culture coverslip to 70% confluency, washed with cold PBS, and fixed with 4% paraformaldehyde in HBSS for 10 min at RT. After washing with HBSS, the cells were permeabilized by incubation with 0.1% Triton X-100 in HBSS at RT for 10 min. After HBSS washing, the coverslip was treated with 3% BSA, 0.2% Tween 20, and 0.2% gelatin in HBSS at RT for 1 h to block non-specific protein adsorption. For Flag-Arc, antibody was diluted at 10 μg/mL in HBSS containing 1% BSA and 1% sucrose at 37 °C for 1 h. The mounting medium for fluorescence with DAPI (Vectashield, Vector Laboratories, Inc., CA, USA) was used for nucleus staining. After being mounted, cells were photographed at x 40 magnification with a fluorescence confocal microscope (LSM510META, Zeiss, Germany).

